# From Structure to Sequence: Identification of polyclonal antibody families using cryoEM

**DOI:** 10.1101/2021.04.13.439712

**Authors:** Aleksandar Antanasijevic, Charles A. Bowman, Robert N. Kirchdoerfer, Christopher A. Cottrell, Gabriel Ozorowski, Amit A. Upadhyay, Kimberly M. Cirelli, Diane G. Carnathan, Chiamaka A. Enemuo, Leigh M. Sewall, Bartek Nogal, Fangzhu Zhao, Bettina Groschel, William R. Schief, Devin Sok, Guido Silvestri, Shane Crotty, Steven E. Bosinger, Andrew B. Ward

## Abstract

One of the rate-limiting steps in analyzing immune responses to vaccines or infections is the isolation and characterization of monoclonal antibodies. Here, we present a hybrid structural and bioinformatic approach to directly assign the heavy and light chains, identify complementarity-determining regions and discover sequences from cryoEM density maps of serum-derived polyclonal antibodies bound to an antigen. When combined with next generation sequencing of immune repertoires we were able to specifically identify clonal family members, synthesize the monoclonal antibodies and confirm that they interact with the antigen in a manner equivalent to the corresponding polyclonal antibodies. This structure-based approach for identification of monoclonal antibodies from polyclonal sera opens new avenues for analysis of immune responses and iterative vaccine design.

**Graphical Abstract:** 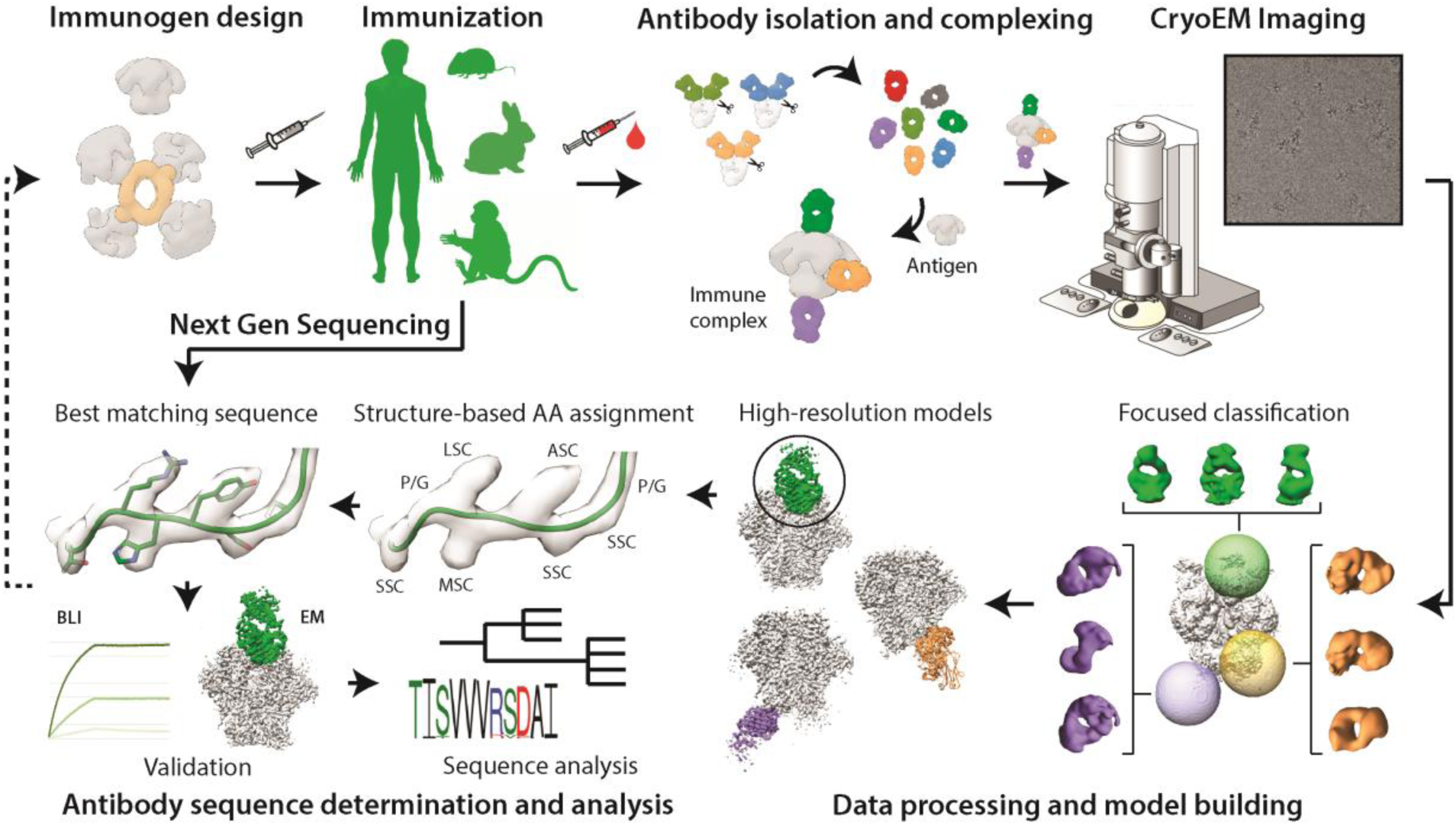

**One Sentence Summary:** CryoEM and next generation sequencing were used to identify monoclonal antibodies elicited by HIV Env vaccine candidates.

## Main Text

Comprehensive analyses of immune responses to infection or vaccination are laborious and expensive (*1, 2*). Classical serology approaches, based on ELISA and viral neutralization assays, offer a wealth of information but require a relatively large set of biological and viral reagents (*3-5*). For rational and structure-based vaccine design efforts it is also necessary to isolate specific monoclonal antibodies (mAbs) elicited by a vaccine/pathogen (*6-11*), yet the isolation of antigen-specific B-cells requires fluorescently labeled probes and access to advanced cell sorting equipment (*12-14*). Individual monoclonal antibodies are subsequently produced and subjected to further binding, structural and functional evaluation to assess epitope specificity, affinity, and activity (i.e. neutralization capacity). High-resolution structural characterization of selected antibodies is most commonly performed at the end of this process, and requires the acquisition of separate datasets for each unique sample (*1, 15, 16*).

Recently, we developed an approach that utilizes electron cryo-microscopy (cryoEM) for characterization of polyclonal antibody (pAb) responses elicited by vaccination or infection (cryoEMPEM), on the level of immune sera (*17*). From a single cryoEMPEM dataset we can readily reconstruct maps of immune complexes at near-atomic resolution (∼3-4 Å range), bypassing the monoclonal antibody isolation steps and streamlining the structural analysis. However, the polyclonal nature of the bound antibodies and the inherent lack of sequence information restricts true atomic resolution for the reconstructed maps, and thereby limits the interpretation of specific epitope-paratope contacts.

Herein, we developed a method to determine monoclonal antibody sequences directly from cryoEMPEM maps. We used structural data from the recently published rhesus macaque immunization experiments with soluble HIV Env trimers (*17, 18*). The three primary cryoEMPEM maps/models used in this study featured BG505 SOSIP bound to structurally distinct polyclonal antibodies (Fig 1). The cryoEM maps were of excellent quality (3.3 - 3.7Å resolution) with high local resolution for the part of the EM map corresponding to the Fab (Fig 1, Fig S1). We used these data and next-generation sequencing (NGS) in a hybrid approach to enable sequence assignment of variable regions of each polyclonal Fab (Fv) including the complementarity determining regions (CDRs).

**Fig 1.**
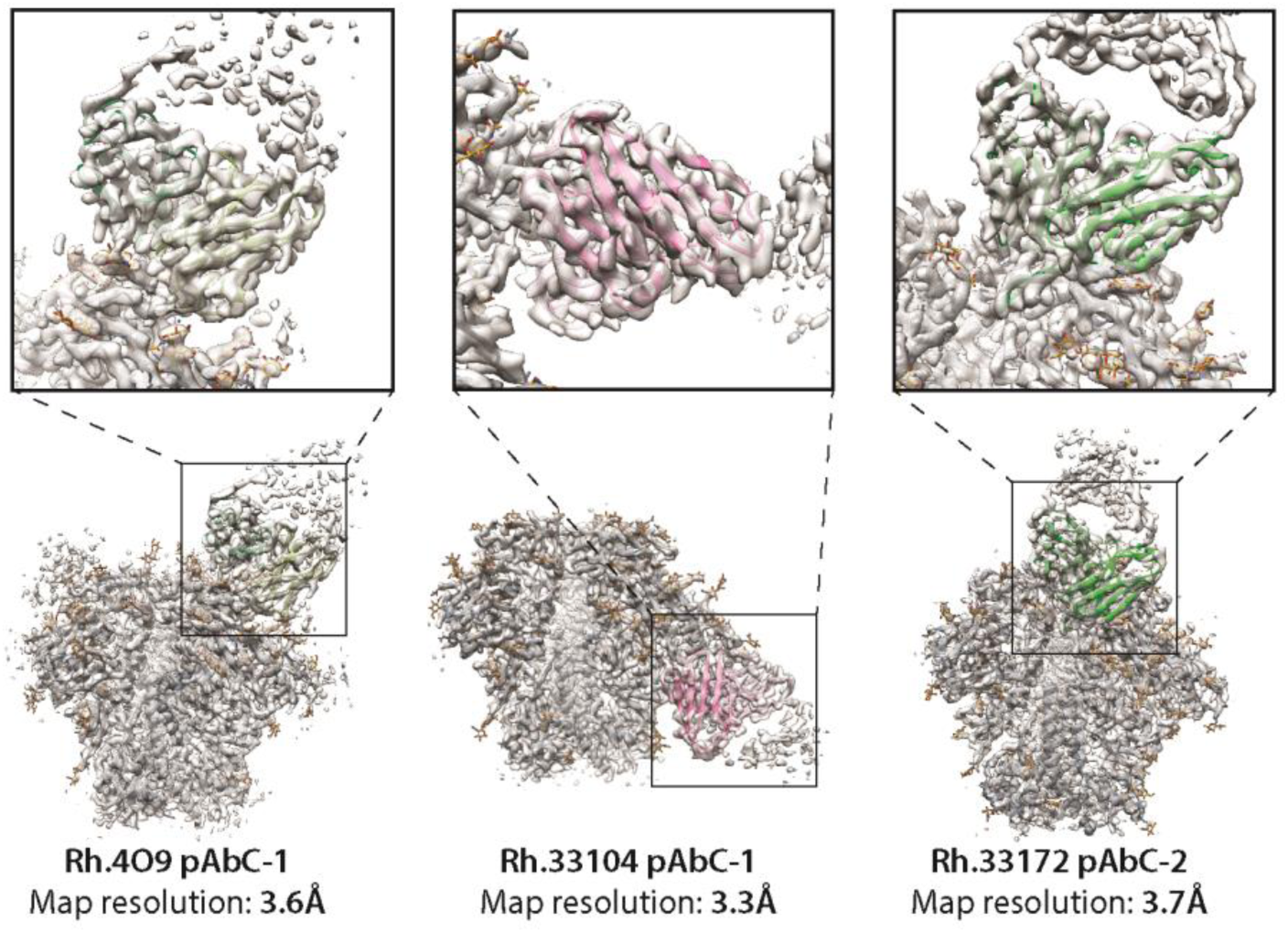
High-resolution maps (transparent gray surface) and corresponding models of immune complexes with polyclonal antibodies (from left to right, Rh.4O9 pAbC-1, Rh.33104 pAbC-1, and Rh.33172 pAbC-2). Full maps and models are shown on the bottom and the close-up view of the epitope-paratope interfaces are shown on top. Ribbon representations of models are used (BG505 SOSIP – dark gray, polyclonal Fab backbone models – green and pink). N-linked glycans are shown as sticks and colored yellow.

First, we analyzed the Rh.4O9 pAbC-1 cryoEMPEM map (Fig 1, left), featuring a polyclonal antibody bound to the V1 loop of BG505 SOSIP antigen. The originally published dataset (*18*) was reprocessed using the focused classification approach shown in Fig S2 to reduce heterogeneity and reconstruct higher-quality map of the V1 pAb. mAbs recognizing this epitope have recently been isolated from the same rhesus macaque (*19*). Two antibodies from the same clonal family, Rh4O9.7 and Rh4O9.8, neutralized the WT BG505 pseudovirus and were found to target the V1 loop through mutagenesis. We applied negative stain electron microscopy (nsEM) to characterize the binding of Rh4O9.8 antibody to BG505 SOSIP and confirmed the V1-specificity (Fig 2A). Furthermore, the Rh4O9.8 mAb superimposed with the polyclonal Fab from the Rh.4O9 pAbC-1 map (Fig 2B).

**Fig 2.**
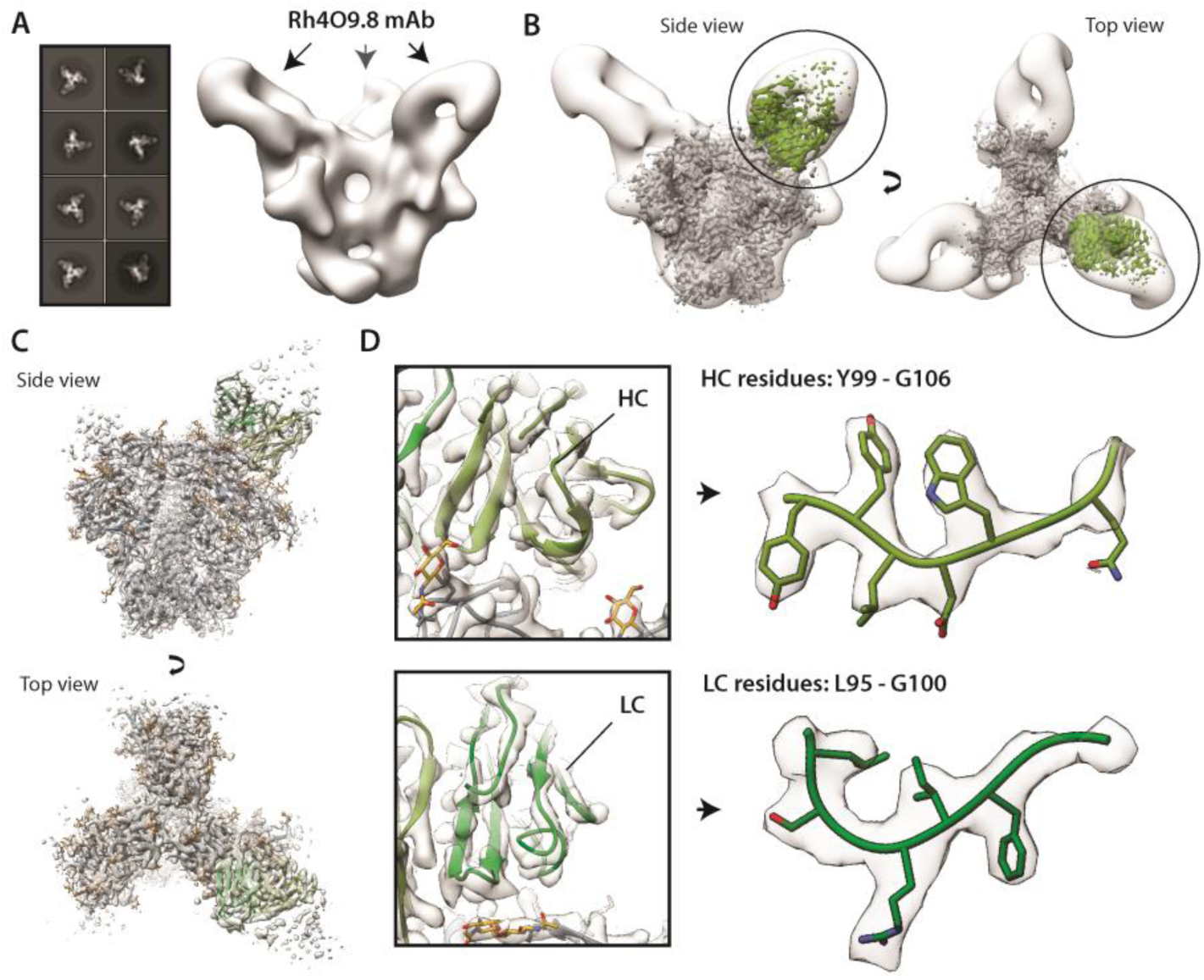
**[A]** nsEM characterization of the Rh4O9.8 Fab in complex with BG505 SOSIP trimer (left – 2D class averages, right – reconstructed 3D map). The arrows show the location of the antibody. **[B]** Overlay of the nsEM map featuring Rh4O9.8 mAb (transparent white surface) and the Rh.4O9 pAbC-1 cryoEMPEM map (BG505 SOSIP - gray; polyclonal antibody - green). **[C]** Ribbon representation of the atomic model of BG505 SOSIP (peptide - dark gray; glycans - yellow) and Rh4O9.8 mAb (green) complex built into the polyclonal cryoEMPEM map, Rh.4O9 pAbC-1 (transparent white surface). **[D]** Close-up view showing model-to-map fit for the heavy chain (top) and the light chain (bottom) of the Rh4O9.8 mAb, including parts of the HCDR3 and LCDR3 (heavy chain – olive green; light chain – forest green; EM map – transparent white surface).

The sequence of Rh4O9.8 was used to build an atomic model into the Fab-corresponding part of the Rh.4O9 pAbC-1 map (Fig 2C). The model exhibited excellent agreement with the cryoEMPEM map at the secondary structure level (Fig 2D, left) as well as the side chain level (Fig 2D, right). Per-residue Q-score plots (*20*) confirm good overall model-to-map fit for the Rh4O9.8 antibody with high degree of consistency between the framework (FW) and the CDR regions (Fig S3). Altogether, the structural data strongly suggest that the Rh4O9.8 mAb isolated by B-cell sorting is a clonal relative of the V1 targeting pAbs identified at the serum level by cryoEM.

In the above example, knowing the sequence of the selected related mAbs enabled interpretation of cryoEM density at an atomic level. However, given the polyclonal nature of bound antibodies and final cryoEMPEM map resolutions of ∼3-4 Å, the structural information is too ambiguous to directly determine antibody sequence from structure alone. We hypothesized that if appropriate sequence databases featuring the BCR repertoire at the corresponding time point were available, the structural information could be used to select the heavy and light chain sequence candidates from those databases. Thus, we developed a hybrid approach that utilizes the structural restraints from cryoEMPEM maps and next generation sequencing (NGS) data of antigen-specific B-cell repertoires for identification of monoclonal antibodies.

First, to approximate the inherent ambiguity in interpreting 3-4 Å resolution cryoEM maps, we developed an assignment system that integrates the degree of certainty associated with the corresponding structural features (i.e. density volume surrounding each amino acid) (Fig 3A, Fig S4). The assignments are performed manually with each amino acid position depicted with a hierarchical category identifier corresponding to a predefined subset of amino acid residues that best correspond to the density. An example of category-based assignment using the Rh.33104 pAbC-1 map is shown in Fig 3B. At the end of this process, structural homology with published antibody structures is used to define the CDR and FW regions, resulting in an initial hierarchical assignment of the antibody sequence.

**Fig 3.**
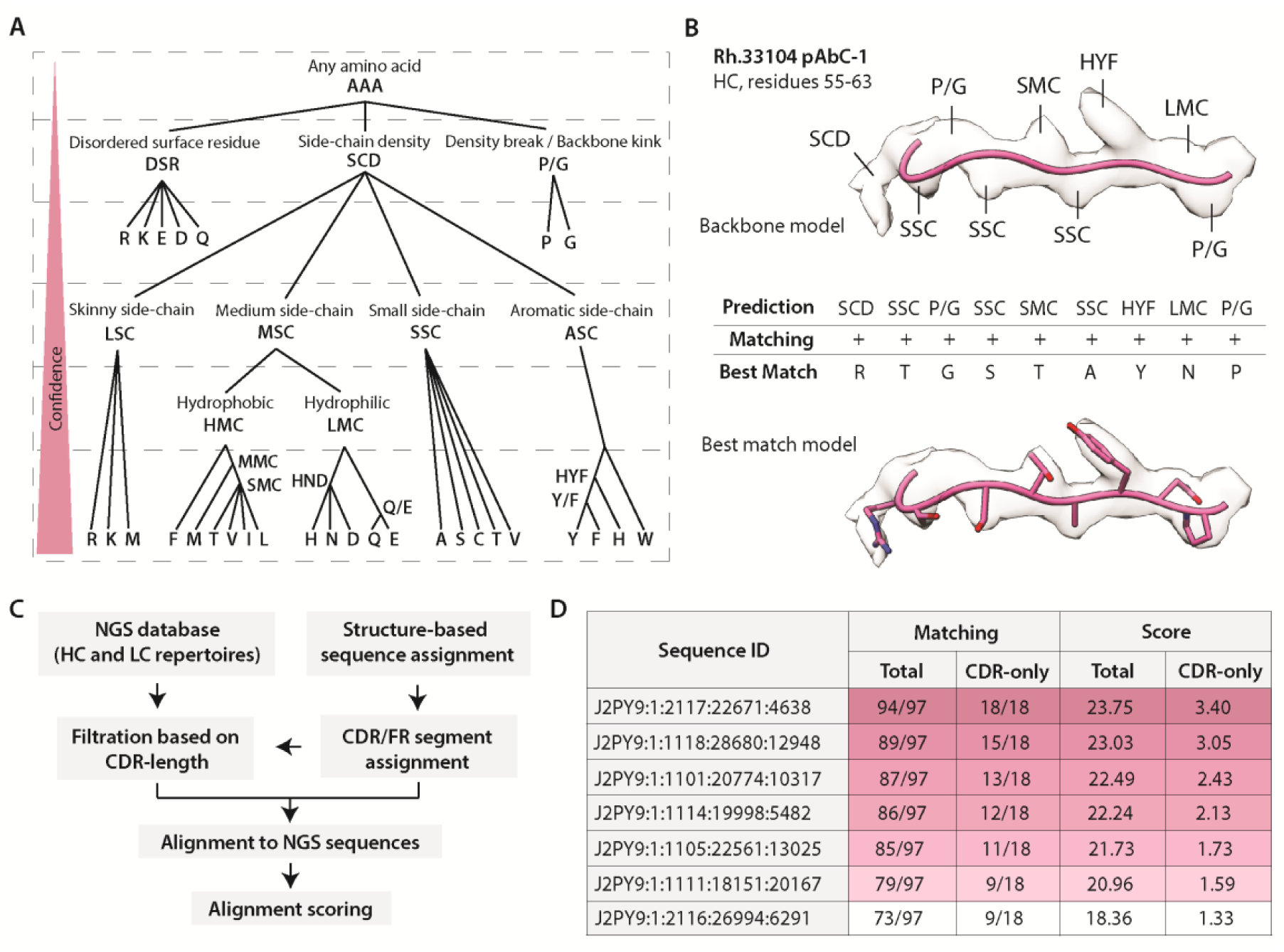
Sequence prediction from high resolution cryoEMPEM maps. **[A]** Amino-acid categories were defined based on common structural features. Hierarchical system allows confidence-based assignment of amino-acids and offers more flexibility in heterogeneous density maps. **[B]** Example of the amino-acid assignment process using Rh.33104 pAbC-1 (heavy chain, residues 55-63). Category assignments are based on the structural features at each amino-acid position. Models are displayed in pink and maps are represented as transparent light-gray surface. The bottom part shows the best matching sequence from the database search and the corresponding model. **[C]** Flowchart of the steps within the sequence alignment and scoring program. **[D]** Examples of matching and scoring results for a subset of NGS sequences, calculated from the search algorithm output data (Rh.33104 dataset, light chain query). The parameters are calculated for the entire aligned portion of the sequence (total) and for CDR segments without the framework (CDR-only). For clarity, the category labels are depicted by 1-3 letter abbreviations in Fig 3A and B. However, in the current version of the program they are numerical (Fig S4).

Next, we designed a search algorithm to take two main inputs (1) preprocessed NGS sequence databases and (2) the user-defined, structure-based sequence queries (Fig 3C). NGS data is pre-filtered by CDR lengths determined during the sequence assignment step by a user-set length tolerance. The program then performs a non-gapped exhaustive alignment search of the query vs every sequence in the database. The queries are split by feature (FW1/2/3, CDR1/2/3) and each feature is aligned independently. Matching is determined based on the agreement of the assigned amino-acid category and the corresponding amino acid from the aligned NGS sequence. For each match (depicted as “+” in the outputs) at a given position, the score is calculated based on the relative ambiguity of the category (1/X_i_ where X_i_ = the number of possible amino acids, X, at position i). Mismatches (depicted as “-” in the outputs) are scored as 0. The output ranks matching sequences based on the CDR lengths, alignment scores and the number and location of mismatches (if any). The calculated score and matching are the two main parameters used to evaluate the agreement of structural data with the sequences from the database (Fig 3D).

To validate the sequence prediction algorithm, we applied it to Rh.33104 pAbC-1 and Rh.33172 pAbC-2 cryoEMPEM datasets (Fig 1). Antibody sequence databases were generated using the germinal center B cells harvested at week 27 time point via fine needle aspiration (*21*); the serum samples used for cryoEMPEM analysis were collected at week 26 and 38 from macaques Rh.33104 and Rh.33172, respectively. Extracted B-cells were sorted based on their binding to the BG505 SOSIP antigen (Fig S5). The sorted B-cells were pooled, lysed and subjected to the next generation sequencing, as described in the methods (See Table S1 for the list of sequencing primers). Notably, the specific heavy and light chain pairing is lost during this process. The total number of sequences recovered for each query are shown in Table S2.

Structure-based sequence category assignments for Rh.33104 pAbC-1 and Rh.33172 pAbC-2 are provided in Auxiliary Supplementary Tables 1 and 2, respectively. The NGS database alignment results with scores are shown in Auxiliary Supplementary Tables 3-6. During the alignment analysis, special emphasis was placed on the score within the CDR regions as the most relevant site for comparisons between different antibodies. The heavy and light chain sequence candidates with the highest scores (for the total sequence and CDRs only) and best matching to predictions were selected and evaluated. Model-to-map fits, alignment and scoring statistics for the best matching Rh.33104 pAbC-1 and Rh.33172 pAbC-2 sequences are shown in Fig S6 and S7, respectively. The mismatches (disagreements between the assigned amino acid category and the amino acid from the best matching sequence) comprised 4 - 18% of the residues in the heavy and light chain sequences (Fig S6B and S7B); for CDRs they occurred in 0 - 14% of cases. Examples of the most common types of mismatches are presented in Fig S8.

Antibodies based on the best matching heavy and light chain sequences from Rh.33104 and Rh.33172 queries were produced and assessed for binding to the BG505 SOSIP antigen using biolayer interferometry (BLI) and sandwich ELISA assays (Fig 4 and Fig S9). Notably, both monoclonal antibodies, termed Rh.33104 mAb.1 and Rh.33172 mAb.1, formed functional dimeric IgG and bound BG505 SOSIP as IgG (Fig 4A,D; Fig S9A,C) and as Fab fragments (Fig S9B,D). EC_50_ values from ELISA experiments with IgGs were 1.93 µg/ml and 2.64 µg/ml and the dissociation constants (K_d_) from BLI were 890 nM and 180 nM, for Rh.33104 mAb.1 and Rh.33172 mAb.1, respectively. These binding affinities are comparable to mAbs isolated from BG505-immunized rhesus macaques in published studies (*6, 19*).

**Fig 4.**
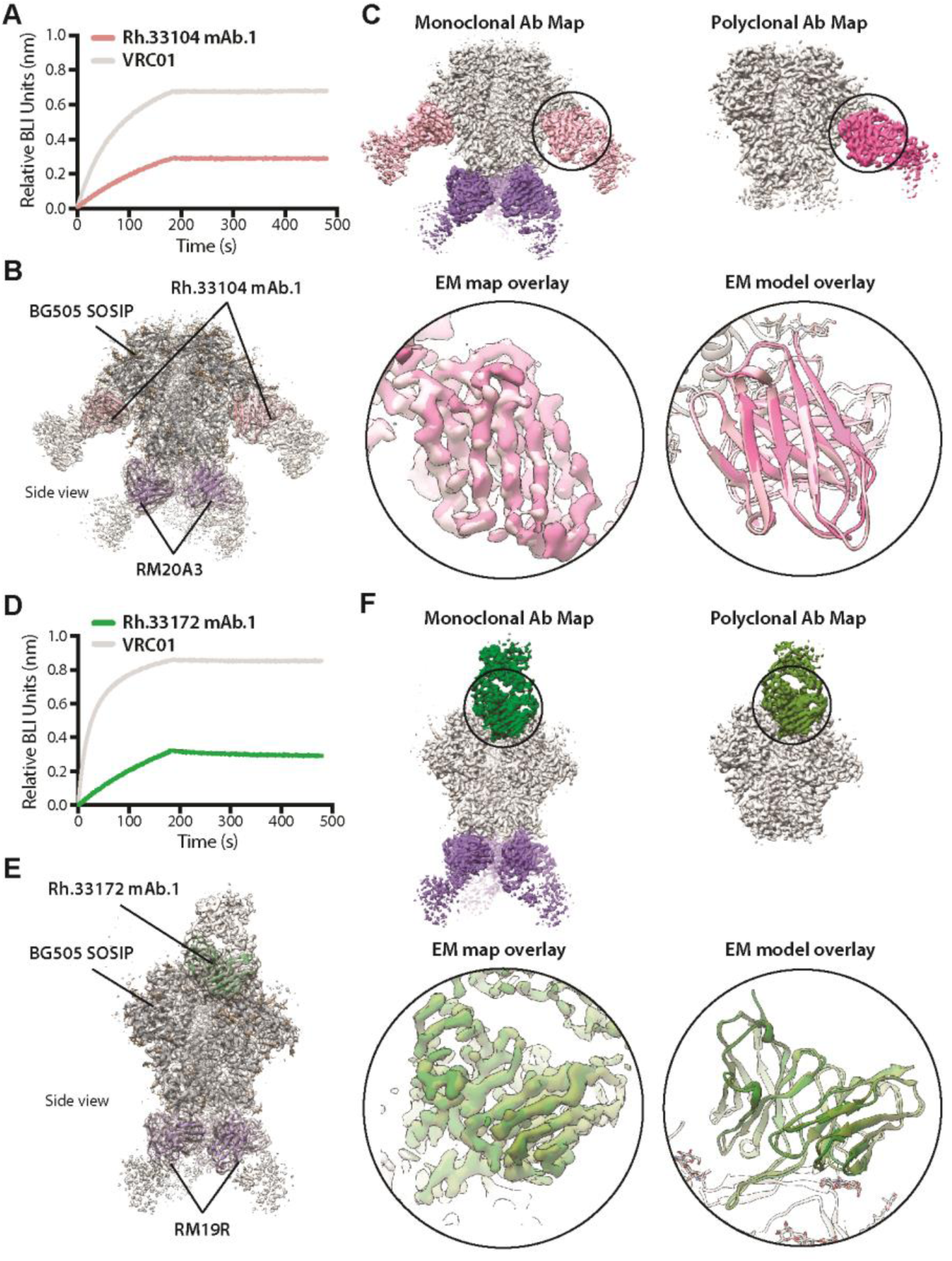
Characterization of the recovered monoclonal antibodies. **[A,D]** Results of the BLI binding analysis performed with Rh.33104 mAb.1 **[A]** or Rh.33172 mAb.1 **[D]** in the form of IgG and the corresponding BG505 SOSIP trimer antigen. VRC01 IgG (gray curves) was included as a positive control and reference. **[B,E]** CryoEM maps and models of the BG505 SOSIP complexes with Rh.33104 mAb.1 **[B]** and Rh.33172 mAb.1 **[E]** as Fab fragments. Maps are represented as transparent gray surface and models are shown as ribbons (BG505 SOSIP – dark gray; RM19R and RM20A3 – purple; Rh.33104 mAb.1 - pink; Rh.33172 mAb.1 – green). **[C,F]** Overlay of the cryoEM maps/models from the EM experiments with monoclonal and polyclonal Fabs for Rh.33104 **[C]** and Rh.33172 **[F]** samples. Full maps of compared immune complexes are shown on top in each panel. Maps are segmented and colored (BG505 SOSIP – light gray; RM20A3 and RM19R – purple; light pink – Rh.33104 mAb.1; deep pink – Rh.33104 pAbC-1; forest green – Rh.33172 mAb.1; olive green – Rh.33172 pAbC-1). Map (left) and model (right) overlays focusing on the Fab are shown in bottom panels. On the backbone level (Cα), the RMSD values between the monoclonal and polyclonal Ab model were 0.668 Å and 0.721 Å for Rh.33104 and Rh.33172 datasets, respectively.

For further validation the two antibodies (as Fabs) were independently complexed with BG505 SOSIP antigen and subjected to cryoEM analysis (Table S3, Fig S10). Monoclonal antibodies targeting the base of the BG505 SOSIP trimer (RM20A3 and RM19R) were used for co-complexing as they help achieve optimal orientation distribution of particles on cryoEM grids. The final map resolutions for the Rh.33104 mAb.1 and Rh.33172 mAb.1 complexes were 3.3 Å and 3.5 Å, respectively. Atomic models were relaxed into the reconstructed maps (Fig 4B,E; Table S4). The structures revealed that both antibodies bound to the same epitopes as the polyclonal antibodies computationally sorted from cryoEMPEM maps (Fig 4C,F). The overlay of the maps/models from experiments with monoclonal and polyclonal Fabs revealed excellent agreement in both cases (bottom panels in Fig 4C,F). Altogether, the structural data confirmed that the monoclonal antibodies selected from the NGS database were clonal members of the polyclonal families detected via cryoEMPEM.

CryoEMPEM is a powerful tool for characterization of polyclonal antibody responses elicited by vaccination or infection. In this study, we expanded the applicability of cryoEMPEM data by introducing a method for identification of functional antibody sequences from structural observations. Conceptually our method is similar to the recently published approach for bottom-up structural proteomics (*22*), in that they both use structural information to infer protein sequences. However, our method utilizes a different assignment system, search algorithm and set of scoring metrics that are specifically optimized for heterogeneous cryoEM density maps, such are the ones obtained by polyclonal epitope mapping.

This approach provides an alternative to traditional mAb discovery methods based on single B-cell sorting, hybridoma and phage display technologies (*12-14*). One of the rate-limiting steps with traditional methods for antibody isolation is screening mAb libraries to identify the clones with desired epitope specificity. Conversely, our approach starts with epitope information for antigen-specific polyclonal antibodies. The structural data is coupled with the corresponding NGS database of antigen-specific BCR sequences, to identify the underlying families of antibodies bound to the epitopes of interest. This effectively circumvents the requirement for single B-cell sorting and monoclonal antibody screening. Therefore, the analysis can be completed within a few weeks of sample collection, instead of a few months; allowing for a more immediate impact on (but not limited to) vaccine design, including real time decision making during immunizations, immunogen redesign for on- and off-target responses, and creation of probes for sorting specific B-cell responses. Importantly, the structure-guided approach eliminates the need for high-resolution characterization of identified monoclonal antibodies as the data is already acquired on the polyclonal level.

Our approach requires high quality structural data (∼4 Å or better map resolutions). Additionally, low-abundance and/or highly diverse classes of antibodies may result in structural information too ambiguous to correctly assign the heavy and light chain sequences. However, novel methods for BCR repertoire determination (e.g. paired heavy-light chain sequencing) and sequence database analysis (e.g. clustering of clonally related sequences), reduce the search space and the relative amount of structural information necessary to identify different polyclonal antibody families. In simpler cases, it may only be necessary to determine the lengths of CDR/FR regions within the heavy and light chain to determine the antibody sequence; this can be readily achieved in maps of intermediate resolution (up to ∼4.5 Å). Further, cryoEM is rapidly improving in terms of throughput and resolution, which should only increase the applicability of our approach.

In this proof-of-concept study, we used lymph node B cells with specificity for the BG505 SOSIP antigen. Our method is anticipated to work with B cells from other sources (e.g., peripheral blood, spleen, bone marrow, plasma cells etc.) and without the pre-sorting for antigen binding. By directly imaging the serum antibodies using cryoEM we have a proxy for abundance, affinity, and clonality. Hence, our approach will open up new doors for both the discovery of monoclonal antibodies as well as analyzing antibody responses to infection and vaccination. The ongoing COVID-19 pandemic has highlighted the need for such robust and rapid technologies.

## Supporting information

Auxilliary Supplemental Table 1

Auxilliary Supplemental Table 2

Auxilliary Supplemental Table 3

Auxilliary Supplemental Table 4

Auxilliary Supplemental Table 5

Auxilliary Supplemental Table 6

## Acknowledgements

The authors express sincere gratitude to Darrell Irvine lab (MIT) and Novavax, Inc. for providing the SMNP and Matrix-M™ adjuvants used in the immunization experiments. The authors also thank Bill Anderson, Hannah L. Turner and Jean-Christophe Ducom (The Scripps Research Institute) for their help with acquisition and processing of electron microscopy data. The authors would also like to acknowledge Lauren Holden for her help with the preparation of the manuscript. This is manuscript number 30087 from The Scripps Research Institute.

## Authors contributions

A.B.W., R.N.K., C.B., C.A.C., G.O., and A.A. conceived the study. D.S., W.R.S., S.C., G.S. and S.E.B. helped with the experimental design. R.N.K. developed the hierarchical assignment system for structure-based sequence inference. C.B. wrote the program for sequence alignment and NGS database search and performed the searches. A.B.W., R.N.K., and C.B. introduced the scoring metrics for alignment of predicted and real sequences. G.O. performed refinement of structural models and helped with the optimization of the sequence identification method. C.A.C. performed pre-processing of NGS sequence data. A.A. performed the sequence assignments for Rh.33104 and Rh.33172 datasets and analyzed the search results. A.A. produced the best matching antibodies and the corresponding antigens and performed the binding and structural analysis. K.C. designed the antigen panel and the sorting protocol. D.G.C. and C.A.E. performed the fine needle extraction of B cells from Rh.33104 and Rh.33172 animals, and B-cell sorting. A.A.U. processed the sorted B cells, subjected them to next-generation sequencing and created the initial sequence databases. L.M.S. and B.N. helped with the acquisition of cryoEMPEM datasets used in this study. B.G. produced the BG505 SOSIP MD39 antigen used for complexing. F.Z. expressed and purified the Rh4O9.8 mAb. A.B.W., S.C., D.S., W.R.S., G.S. and S.E.B. supervised the study and provided essential guidance. A.A., C.B., and A.B.W. wrote the original draft of the manuscript. All authors contributed to the manuscript text by assisting in writing and/or providing critical feedback.

## Data and materials availability

The sequence alignment program will be made available upon request. It will also be released on GitHub (https://github.com/) following the publication of this manuscript. 3D maps and models from the electron microscopy experiments have been deposited to the Electron Microscopy Databank (http://www.emdatabank.org/) and the Protein Data Bank (http://www.rcsb.org/), respectively. The accession numbers are listed in the table below:

| Sample description | EMDB ID | PDB ID |
| --- | --- | --- |
| Negative stain EM map of BG505 SOSIP in complex with Rh4O9.8 Fabs | EMD-23778 | N/A |
| Rh.4O9 pAbC-1 cryoEMPEM map (model features BG505 SOSIP in complex with Rh4O9.8 Fab) | EMD-23779 | 7MDT |
| cryoEM map and model of the BG505 SOSIP / Rh.33104 mAb.1 / RM20A3 complex | EMD-23780 | 7MDU |
| cryoEM map and model of the BG505 SOSIP / Rh.33172 mAb.1 / RM19R complex | EMD-23801 | 7MEP |

