## Supplemental Materials for "From Structure to Sequence: Identification of polyclonal antibody families using cryoEM"

#### Methods

**- Antigen expression and purification**

Antigen expression and purification were performed as described previously (*1*). Briefly, BG505-SOSIP.v5.2(7S) N241/N289 (subcloned into a pPPI4 vector) and BG505 SOSIP MD39 (subcloned into a pHLsec vector) construct genes were expressed in 293F cells (Thermo Fisher Scientific). The proteins were purified from cell supernatants using PGT145 or 2G12 immunoaffinity chromatography. 3M $MgC1_2$ buffer was used for protein elution from the immunoaffinity matrix. BG505 SOSIP samples were concentrated, buffer exchanged to TBS (Alfa Aesar) and subjected to size-exclusion chromatography (SEC). HiLoad 16/600 Superdex 200 pg (GE Healthcare) running TBS buffer was used for SEC purification. Fractions corresponding to the BG505 SOSIP antigen were pooled, concentrated to 1 mg/ml and frozen for storage.

**- Rhesus macaque immunizations**

The rhesus macaque immunization experiments have been reported previously (*1*). Immunogens were administered subcutaneously, divided between the right and left mid-thighs at weeks 0, 8, 24 and 36. Animals were immunized with BG505 SOSIP MD39 trimer (100 μg per dose) or BG505 SOSIP T33-31 nanoparticle (119 μg per dose) with Matrix-M$^{TM}$ (Novavax; 75 μg per dose) or SMNP (Darrell Irvine Lab, MIT; 750 U per dose) adjuvants. Blood draws were performed biweekly. Lymph node fine needle aspirates (FNA) were performed as previously described (*2*) at weeks 8, 11, 14, and 27. Lymph node biopsies were performed between weeks 40 and 42. Animal work was performed at the Yerkes National Primate Research Center, Atlanta, GA, USA. All procedures were approved by Emory University Institutional Animal Care and Use Committee protocol 201700723. Animal care facilities are accredited by the U.S. Department of Agriculture and the Association for Assessment and Accreditation of Laboratory Animal Care International.

For structural cryoEMPEM analyses of polyclonal antibody responses in animals Rh.33104 and Rh.33172, we used the serum samples from week 26 and week 38, respectively (*1*). The reconstructed EM maps and models correspond to these time points. B-cell repertoire databases were generated using the FNA samples from week 27.

**- B-cell sorting**

Biotinylated BG505 SOSIP MD39 trimer and BG505 SOSIP.v5.2 N241/N289 trimers were generated as previously described (*1, 2*). Biotinylated proteins were individually premixed with fluorochrome-conjugated streptavidin (Brilliant Violet 650 or Brilliant Violet 421, BioLegend) at RT for 20 minutes. BG505 SOSIP MD39-ferritin and BG505 SOSIP T33-31 nanoparticles were generated and directly conjugated to Alexa Fluor 647 (Thermo Fisher). Cells were incubated with indicated probes for 30 minutes at 4°C, and then surface antibodies were added and incubated for 30 minutes at 4°C. Cells were washed and then sorted on a FACSAria II. The sorting procedure is shown in Fig S5.

Rh.33104 (BG505 SOSIP MD39 immunized) – LN cells collected at week 27 were stained with the following probes and antibodies: BG505 SOSIP MD39 trimer-Brilliant Violet 650, BG505 SOSIP MD39 trimer- Brilliant Violet 421, BG505 SOSIP MD39-ferritin nanoparticle-Alexa Fluor 647, fixable viability dye-eFluor 780 (Thermo Fisher), mouse anti- human CD4 APC eFluor 780 (SK3, Thermo Fisher), mouse anti-human CD8 APC eFluor 780 (RPA-T8, Thermo Fisher), mouse anti-human CD16 APC eFluor 780 (ebioCB16, Thermo Fisher), mouse anti-human CD20 Alexa Fluor 488 (2H7, BioLegend), mouse anti-human IgG PE-Cy7 (G18-145, BD Biosciences), mouse anti-human IgM PerCP-Cy5.5 (G20-127, BD Biosciences), mouse anti-human CD38 PE (OKT10, NHP Reagent Resource), mouse anti-human CD71 PE-CF594 (L01.1, BD Biosciences).

Rh.33172 (BG505 SOSIP T33-31 nanoparticle immunized) – LN cells collected at week 27 were stained with the following probes and antibodies: BG505 SOSIP MD39 trimer-Brilliant Violet 650, BG505 SOSIP.v5.2 N241/N289 trimer-Brilliant Violet 421, BG505 SOSIP T33-31 nanoparticle-Alexa Fluor 647, fixable viability dye-eFluor 780 (Thermo Fisher), mouse anti-human CD4 APC eFluor 780 (SK3, Thermo Fisher), mouse anti-human CD8 APC eFluor 780 (RPA-T8, Thermo Fisher), mouse anti-human CD16 APC eFluor 780 (ebioCB16, Thermo Fisher), mouse anti-human CD20 Alexa Fluor 488 (2H7, BioLegend), mouse anti-human IgG PE-Cy7 (G18-145, BD Biosciences), mouse anti-human IgM PerCP-Cy5.5 (G20-127, BD Biosciences), mouse anti-human CD38 PE (OKT10, NHP Reagent Resource), mouse anti-human CD71 PE-CF594 (L01.1, BD Biosciences).

**- NHP Immunoglobulin Repertoire Library Preparation and Sequencing**

IG repertoire sequencing libraries were prepared using a protocol provided by Dr. Daniel Douek, NIAID/VRC (*3*) and similar to described previously (*2*). Primer sequences are listed in Table S1. With the exception of the Oligo dT and SMARTer II A Oligo template-switch primer, all oligos were obtained from Integrated DNA Technologies. Qiagen RNeasy kits (Valencia, CA) were used to isolate RNA from isolated antigen-specific B cells obtained from LN FNAs at the post-vaccine time points described above. cDNA was generated using Clontech SMARTer kits: 8 µL RNA was mixed with 1 µL of 5' CDS oligo(dT) (12 µM) and incubated at 72˚C for 3 minutes and 4˚C for 1 minute. Following the addition of 8.5 µL RT mastermix comprised of 5x RT Buffer [250 mM Tris-HCl (pH 8.3), 375 mM KCl, 30 mM MgCl2], Dithiothreitol, DTT (20 mM), dNTP Mix (10 mM), RNAse Out (40U/µL), SMARTer II A Oligo (12 µM) and Superscript II RT (200U/µL)), samples were incubated for at 42˚C for 90 minutes and 70˚C for 10 minutes. AMPure XP beads (catalog# A63882) were used for purifying first-strand cDNA. To amplify IgG, IgK or IgL variable regions from cDNA, we used the KAPA Real Time Library Amplification Kit (catalog# KK2702). The master mix was made up of 2X KAPA PCR Master Mix, 12 µM µL 5PIIA and 5 µL of IgG, IgK or IgL Constant Region Primers (2 µM) (*4*). 19.3 µL of cDNA and 30.7 µL of master mix were mixed followed by centrifugation at 2000 RCF for 1 minute. This was followed by monitoring the real-time PCR. The PCR was stopped in the exponential phase (23 cycles for IgG and ~18 cycles for IgK/IgL) and AMPure XP beads were used for purification of the amplified products. Barcodes and Illumina adapters were added using two subsequent rounds of PCR. In the first step (addition of barcodes) grafting primers (forward P5_Seq BC_XX 5PIIA, 1 µL of 10 µM stock) (reverse P7_i7_XX_Ig, 1 µL of 10 µM stock) containing randomized stretch of 4-8 random nucleotides and unique barcodes were mixed with 46 µL of master mix (2X KAPA PCR Master Mix 2x, SYBR Green 1:10K, Nuclease-free water) and 2 µL of 1:10 diluted IgG/IgK/IgL amplicon; mixtures were amplified using real-time PCR (7 cycles for IgG and 6 cycles for IgK/IgL) and purification of the library using AMPure XP Beads. The final step involved the grafting of Illumina adapters for which 34 µL of master mix (2X KAPA PCR Master Mix, 10 µM P5_Graft P5_seq, Nuclease-free water) was mixed with 1 µL of 10 µM P7_I7_XX Ig constant region primer and 15 µL of purified product from the previous, and amplification by real-time PCR (6 cycles) and final purification of the library with AMPure XP beads. Agilent Bioanalyzer was used to assess the quality of the libraries. The libraries were then pooled, and the sequencing was carried out on an Illumina MiSeq (309 paired-end).

**- NGS data processing and sequence analysis.**
Read pairs were assembled and filtered for length and quality using the pRESTO toolkit (*5*). Sequence regions corresponding to the primers were masked and duplicate sequences were collapsed (*5*). Germline antibody gene assignment was conducted with IgBLAST using a comprehensive Indian origin RM germline BCR database (*6, 7*). IgBLAST results were filtered for productive sequences and sorted based on feature lengths. For heavy chain queries a single NGS database is used for each antibody. For light chains, the search is performed independently for the Ig-κ and Ig-λ databases.

**- cryoEMPEM maps and models used in the analysis**
Rh.4O9 pAbC-1. The Rh.4O9 pAbC-1 map was generated by applying the focused classification approach (*1*) to the Rh.4O9 cryoEMPEM dataset that was published previously (*8*). This data was reprocessed to resolve a map of higher quality (i.e. higher local resolution for the Fab). The data processing workflow is illustrated in Fig S2. The resulting EM map for Rh.4O9 pAbC-1 (Fig 1 and Fig S1) has been uploaded to the Electron Microscopy Data Bank (EMDB), entry ID: EMD-23779. The structural model of the complex, comprising BG505 SOSIP.v5.2 (*9*) and Rh4O9.8 mAb (*10*), was built into the EM map. ABodyBuilder (*11*) was applied to create an initial model of the Fab. Previously published structure of BG505 SOSIP (PDB ID: 5CEZ (*12*)), was used as a starting model for the antigen. The final model of the complex was generated by applying iterative rounds of manual refinement in Coot (*13*) and automated refinement in Rosetta (*14*). For validation we applied EMRinger (*15*) and MolProbity (*16*) software packages (Table S4). Per-residue characterization of the model to map fit (shown in Fig S3) was performed in UCSF Chimera (*17*) using the Q-score plugin (*18*). The refined model of the complex was submitted to the Protein Data Bank (PDB), entry ID: 7MDT. HxB2 numbering was used for the BG505 SOSIP antigen and Kabat numbering was applied for Rh4O9.8 antibody.

Rh.33104 pAbC-1 and Rh.33172 pAbC-2. The assignment of polyclonal antibody sequences was performed using previously published cryoEMPEM data (*1*). Specifically, for Rh.33104 pAbC-1 and Rh.33172 pAbC-2, we used the maps uploaded under EMDB IDs: EMD-23227 and EMD-23232, respectively. The corresponding structures were previously uploaded under PDB IDs: 7L8A and 7L8F, respectively. In the structures, the polypeptide backbone of each antibody was represented as a poly-alanine pseudo-model. The number of amino-acid residues was adjusted to achieve the most optimal model-to-map fit on the backbone level.

**- Antibody sequence assignment**
Sequence assignment the heavy and light chains was performed manually in Coot, using the models and maps from the Rh.33104 pAbC-1 and Rh.33172 pAbC-2 datasets. We developed a system for amino-acid assignment based on the corresponding structural features, that also takes into consideration the degree of certainty associated with each assignment. Density volume surrounding each amino acid is attributed a hierarchical category identifier representing a predefined subset of amino acid residues that best correspond to the density. The category assignment tree is shown in Fig S4. In the final output, the heavy and light chain sequences are represented as strings of numerical category identifiers (for examples see Auxiliary Supplementary

Tables 1. and 2.). In addition to shape properties, our system allows to further categorize certain amino acid groups (e.g. medium side-chain group) based on the local environment (i.e. hydrophobic/hydrophilic) which helps narrow down the list of possible amino acids. Structural homology with published rhesus macaque antibody structures (PDB IDs: 4KTE, 4KTD, 4RFE, 4Q2Z (*19-21*)) is applied at the end of the assignment process to define the CDR and FW regions within the antibody.

**- Sequence alignment and scoring**

Sequence match searching was performed using Python 3.6.3 (www.python.org) in a Jupyter Notebook (www.jupyter.com) environment. To prepare for alignment, the hierarchical ambiguity codes from Fig 3A were first translated into numerical codes and organized into a recursive binary tree. This allows the algorithm to call upon any specific branch and fetch all downstream amino acid possibilities in an efficient manner. The search can be performed on two main data formats: protein FASTA files and .tsv formatted output from igblastn results. The search can also compare the ambiguity codes directly to the allele database present in the Indian Origin Rhesus macaque Germ Line Database (GLD) currently hosted by the Ward lab. (ward.scripps.edu/gld; (*7*)). Search results against FASTA files and the GLD are basic and do not return information outside of alignment of the main section of sequence. Searches run on .tsv files from IGBLASTn (referenced below as “data frame”) allow the user to pass queries in a split method based on IMGT designations for relevant framework and CDR loop regions (*6, 22*). For the main searches featured here, query strings of ambiguity codes were manually segmented into the following regions: FW1, CDR1, FW2, CDR2, FW3, CDR3. Before searching, sequences flagged as unproductive by igblastn are removed and the data frame to be searched is subjected to filtering based on user-defined criteria. The user can filter out entries from the data frame based on any column that exists, typically restricting based on a range of desired lengths of FW and CDR regions. Once the data frame is prepared and a search is initiated, the algorithm exhaustively compares each query segment to the relevant column from the data frame in pairwise fashion. Alignment is allowed to shift by a factor defined by the user (default 2 AA), and results are returned based on a maximum alignment score for each query to each subject sequence. Scores are calculated as the inverse of the number of amino acids represented by each ambiguity code at each position, but only if there is a match. For example, if an ambiguity code represents a branch that has 5 possible amino acids and there is a match, the position is assessed a score of 1/5, or 0.2. Scores range from 1/20 (0.05) to 1/1 (1.00). Scores are tallied for each query section of each row of the data frame and returned to the user in csv format for easy manipulation and selection of high scoring alignments. The sequence alignment program is available upon request. It will also be released on GitHub (https://github.com/) following the publication of this manuscript.

**- Analysis of sequence alignment results**

For search result analysis, we calculated the alignment scores for the entire sequence (Total Score) and the complementarity-determining regions (CDR-only Score). The heavy and light chain sequences featuring a combination of the highest Total and CDR-only scores from each search were selected for subcloning and expression. For analysis of the light chain alignment results, we have also calculated the mean total alignment scores for all NGS sequences in the corresponding Ig-κ and Ig-λ queries. The comparison of maximum and mean total alignment scores from the two queries was applied to determine if the antibody light chain was Ig-κ or Ig-λ. For sequences and alignment scores see Auxiliary Supplementary Tables 3-6.

**- Antibody expression and purification**

Top-scoring heavy and light chain sequences from the corresponding NGS database searches were subcloned into the AbVec-hIgG1 and AbVec-hIgKappa expression vectors, respectively (*23, 24*). Sanger sequencing was applied to verify the final DNA vectors. 500μg of the heavy chain and 250 μg of the light chain DNA expression vectors were applied for co-transfection of 1L of HEK293F cells to produce Rh.33104 mAb.1 and Rh.33172 mAb.1 (as full IgG and Fab fragments). PEI MAX (Polysciences, Inc.) was used as a transfection reagent at three-fold mass excess with respect to total DNA amount. Antibodies were purified from cell supernatants using the MAbSelect Xtra (GE Healthcare Life Sciences) and CaptureSelect IgG-CH1 (ThermoFisher Scientific) columns for IgG and Fab purification, respectively. Antibody samples were concentrated, buffer exchanged to TBS buffer (Alfa Aesar), and then subjected to SEC purification (HiLoad 16/600 Superdex S200 pg column; GE Healthcare Life Sciences).

**- Negative stain EM of BG505 SOSIP in complex with Rh4O9.8 mAb**

Rh4O9.8 monoclonal antibody (*10*) was kindly provided by the Devin Sok lab (International AIDS Vaccine Initiative - Neutralizing Antibody Center, La Jolla, CA, USA). 15μg of BG505 SOSIP MD39 was incubated with 15μg of the Rh4O9.8 antibody (as Fab fragment) for 1 hour at room temperature. The complex was then subjected to SEC purification (Superose 6 Increase column, 10/300 GL, GE healthcare) with TBS (10mM Tris-HCl, 150mM NaCl, pH 7.4) as the running buffer. Fractions corresponding to the immune complex were pooled and concentrated using an Amicon filter unit with 10 kDa cutoff (EMD Millipore). For negative stain EM, the SOSIP-Fab complex was diluted to 20 μg/ml, and 3μl were applied onto a carbon-coated copper grid (400-mesh, Electron Microscopy Sciences). The grid was pre-treated by glow-discharge for 30 s. The complex solution was blotted-off after 10s and the grid was subsequently stained with 2% (w/v) uranyl formate for 60 s. Imaging was performed on a Tecnai F20 electron microscope operating at 200keV (1.77 Å/pixel; 62,000X magnification) as described previously (*25*). The defocus was set to −1.50 μm and the electron dose was adjusted to 25 $e^-/Å^2$ using Leginon software (*26*). All 2D and 3D classification and 3D refinement steps were conducted in Relion 3.0 (*27*). EM density maps were visualized in UCSF Chimera (*17*). Reconstructed EM map of BG505 SOSIP in complex with 3 Rh4O9.8 Fabs was submitted to Electron Microscopy Data Bank (EMDB ID: EMD-23778).

**- Sandwich ELISA assays**

Sandwich ELISA experiments were performed with Rh.33104 mAb.1 and Rh.33172 mAb.1 (as IgG). All experiments were performed in triplicates. BioStack Microplate Stacker system (BioTek) was used for buffer addition and wash steps. 12N antibody with specificity towards the base of BG505 SOSIP, was diluted to 3μg/ml and immobilized onto high-binding, 96-well microplates (Greiner Bio-One) for 2 hours at room temperature. The plates were washed 3 times with TBST buffer (TBS + 0.1% Tween-20) and blocked overnight with TBS + 5% bovine serum albumin (BSA) + 0.05% Tween-20 at 4°C. Plates were washed 3 times with TBST, followed by the addition of the antigen solution (PBS + 1% BSA + 3 μg/ml of BG505 SOSIP). For Rh.33172 mAb.1 experiments we used BG505 SOSIP.v5.2(7S) N241/N289 construct; for Rh.33104 mAb.1 experiments we used BG505 SOSIP MD39. This was done to match the BG505 SOSIP construct to the original immunogen that elicited the corresponding polyclonal antibodies in rhesus macaques (*1*). The plates were incubated with antigen solution for 2 hours and then washed 3 times with TBST. Serial 3-fold dilutions of Rh.33104 mAb.1 or Rh.33172 mAb.1 (starting at 100 μg/ml) were prepared in TBS and added to plates coated with corresponding antigen. The plates were incubated for 2 hours at room temperature and subsequently washed 3 times with TBST. AP-conjugated AffiniPure goat anti-human IgG, (Jackson Immunoresearch, Cat # 109-055-097) was diluted 1:4000 in TBS + 1% BSA buffer and added to each well for 1 hour at room temperature. Following 3 wash steps with TBST, 1-Step PNPP Substrate Solution (Thermo-Fisher Scientific) was applied to each well for detection. Synergy H1 plate reader (BioTek) was used for acquisition of colorimetric data by recording the absorbance at 405 nm wavelength. Data was analyzed in Graphpad Prism (version 8.4.3) software and midpoint titers ($EC_{50}$) were determined.

**- Biolayer interferometry (BLI)**

Octet Red96 instrument (ForteBio) was used for BLI data collection. Antibody and antigen solutions were prepared in kinetics buffer (DPBS + 0.1% [w/v] BSA + 0.02% [v/v] Tween-20). All BLI experiments were conducted at 25 °C. BLI experiments with IgGs were performed as described previously (*1, 28*). Rh.33104 mAb.1, Rh.33172 mAb.1 and VRC01 antibodies (as IgGs) were diluted to 5 µg/ml and immobilized onto human IgG Fc capture (AHC) biosensors (ForteBio). VRC01 served as a positive control. Antibody-coated sensors were then transferred to wells with corresponding BG505 SOSIP antigens (see ELISA method section above for explanation) at 1000 nM concentration. Association and dissociation steps were set to 180 s and 300 s, respectively. Data was processed using Octet System Data Analysis v9.0 (ForteBio). Negative control measurements (with kinetics buffer) were used for background correction. Final plots were prepared in Graphpad Prism (version 8.4.3).

Experiments with Fab fragments were performed as described previously (*7*). Fabs were diluted to 25µg/ml and immobilized onto anti-human Fab-CH1 (FAB2G) biosensors (ForteBio). Serial 2-fold dilutions of the corresponding BG505 SOSIP antigens (see ELISA method section above for explanation) were prepared for binding studies, starting at 2000nM. The lengths of association and dissociation steps were set to 600 and 1200s, respectively. Data processing and determination of kinetic parameters were performed in Octet System Data Analysis v9.0 software (ForteBio). Data plots were prepared in Graphpad Prism (version 8.4.3).

**- CryoEM analysis of monoclonal antibody complexes – Preparation of complexes.**

Rh.33104 mAb.1 complex preparation: 250 µg of BG505 SOSIP MD39 was incubated with 600 µg of Rh.33104 mAb.1 Fab and 600 µg of RM20A3 Fab (*7, 29*), at room temperature, overnight. The complex was SEC-purified using a HiLoad 16/600 Superdex pg200 (GE Healthcare) column, with TBS as a running buffer. SEC fractions corresponding to the complex were pooled and concentrated to 6.0 mg/ml using an Amicon filter unit with 10 kDa cutoff (EMD Millipore).

Rh.33172 mAb.1 complex preparation: 250 µg of BG505 SOSIP.v5.2(7S) N241/N289 was incubated with 600 µg of Rh.33172 mAb.1 Fab and 600 µg of RM19R Fab (*7*), at room temperature, overnight. All other purification steps were equivalent as with Rh.33104 mAb.1 complex.

Antibodies RM20A3 and RM19R, with specificity towards the trimer base, were used for co-complexing because in the past we have observed that they can improve the orientational distribution of HIV Env trimers on cryoEM grids (*29*).

**- CryoEM analysis of monoclonal antibody complexes – Grid preparation.**

UltrAuFoil R 1.2/1.3 grids (Au, 300-mesh; Quantifoil Micro Tools GmbH) were used for sample vitrification. The grids were treated with $Ar/O_2$ plasma (Solarus 950 plasma cleaner, Gatan) for

10s immediately prior to sample application. 0.5 µL of 0.04 mM lauryl maltose neopentyl glycol (LMNG) stock solution was mixed with 3.5 µL of the complex and 3 µl were immediately loaded onto the grid. Grids were prepared using Vitrobot mark IV (Thermo Fisher Scientific). Temperature inside the chamber was maintained at 10˚C while humidity was at 100%. Blotting force was set to 0, wait-time to 10 s while the blotting time was varied within a 4-7 s range. Following the blotting step, the grids were plunge-frozen into liquid ethane, cooled by liquid nitrogen.

**- CryoEM analysis of monoclonal antibody complexes – Data collection and processing.**
Samples were imaged on FEI Titan Krios electron microscope (ThermoFisher) operating at 300 keV. The microscope was equipped with the K2 summit detector (Gatan) operating in counting mode. Exposure magnification was 29,000 and the pixel size was 1.03 Å (at the specimen plane). Leginon software suite (*26*) was used for automated data collection. Data collection information for the two datasets featuring different monoclonal antibody complexes are presented in Table S3. Micrograph movie frames were aligned and dose-weighted using MotionCor2 (*30*) and GCTF (*31*) was applied for estimation of CTF parameters. Initial processing steps (particle picking and 2D classification) were performed in cryoSPARC.v2 (*32*). Ab initio refinement in cryoSPARC was applied to generate the initial reference model for each complex. Clean particle stack was subsequently transferred to Relion/3.0 (*27*) for further 2D and 3D processing steps. Data processing workflows and relevant information are presented in Fig S10.

**- CryoEM analysis of monoclonal antibody complexes – Model building and refinement**
Postprocessed cryoEM maps from Relion/3.0 were used to generate atomic models. PDB entry 6vfl (*33*) was used as initial model for BG505 SOSIP-corresponding part of the complex. The sequence was adjusted to match the exact BG505 SOSIP variant used for the preparation of imaged monoclonal antibody complex (BG505 SOSIP MD39 or BG505 SOSIP.v5.2(7S) N241/N289). Initial models for RM20A3 and RM19R antibodies were adapted from PDB 6X9R (*29*) and PDB 6VKN (*7*), respectively. ABodyBuilder (*11*) was applied to create the initial Fab models for Rh.33104 mAb.1 and Rh.33172 mAb.1. Individual components were docked into the corresponding parts of each cryoEM map in UCSF Chimera (*17*) to create the initial atomic models. Iterative rounds of manual model refinement in Coot (*13*) and automated refinement using Rosetta (*14*) were performed to produce the final models. HxB2 numbering was used for BG505 SOSIP antigens and Kabat numbering was applied for antibodies in each complex. For model validation we applied EMRinger (*15*) and MolProbity (*16*) packages. Model refinement statistics is reported in Table S4. The refined models were submitted to the Protein Data Bank (PDB ID: 7MDU, 7MEP).

### Supplementary Figures and Tables

**Auxiliary Supplementary Table 1**. Structure-based sequence assignment for Rh.33104 pAbC-1 (provided externally)
**Auxiliary Supplementary Table 2.** Structure-based sequence assignment for Rh.33172 pAbC-2 (provided externally)
**Auxiliary Supplementary Table 3.** Search results for Rh.33104 light chain (Ig-κ) dataset (provided externally)
**Auxiliary Supplementary Table 4.** Search results for Rh.33104 heavy chain dataset (provided externally)
**Auxiliary Supplementary Table 5.** Search results for Rh.33172 light chain (Ig-κ) dataset (provided externally)
**Auxiliary Supplementary Table 6.** Search results for Rh.33172 heavy chain dataset (provided externally)

**Table S1.** Primer sequences used for NHP IgG/IgK/IgL repertoire sequencing.

| Primer | Sequence |
|---|---|
| **CDS Oligo dT** | TTTTTTTTTTTTTTTTTTTTTTTTTVN |
| **SMARTer II A Oligo** | AAGCAGTGGTATCAACGCAGAGTACATrGrGrG |
| **IgG** | CCAGGGGGAAGACCGATGGGCCCTTGGTGGA |
| **IgK** | GCGGGAAGATGAAGACAGATGGTGCAGCCACAG |
| **IgL** | GGCCTTGTTGGCTTGAAGCTCCTCAGAGGAGGG |
| **P5_Seq BC_XX 5PIIA** | CACGACGCTCTTCCGATCT NNNN AACCACTA AAGCAGTGGTATCAACGCAGAGT |
| **P7_ i7_XX IgG** | CAAGCAGAAGACGGCATACGAGAT TAGTGGTT GCCAGGGGGAAGACCGATGGGCCCTTGGTGGA |
| **P7_ i7_XX IgK** | CAAGCAGAAGACGGCATACGAGAT TAGTGGTT GCGGGAAGATGAAGACAGATGGTGCAGCCACAG |
| **P7_ i7_XX IgL** | CAAGCAGAAGACGGCATACGAGAT TAGTGGTT GGCCTTGTTGGCTTGAAGCTCCTCAGAGGAGGG |
| **P5_Graft P5_seq** | AATGATACGGCGACCACCGAGATCTACAC TCTTTCCCTACACGACGCTCTTCCGATCT |

**Table S2.** Sequence count for different NGS datasets

| Sequence count | Rh.33104 Ig-κ | Rh.33104 Ig-λ* | Rh.33104 IgH | Rh.33172 Ig-κ | Rh.33172 Ig-λ* | Rh.33172 IgH |
|---|---|---|---|---|---|---|
| **Starting NGS dataset** | 178299 | 209168 | 137916 | 192252 | 235499 | 197099 |
| **Filtered by CDR lengths** | 70845 | 42185 | 5578 | 117254 | 111770 | 4428 |
| **Selected** | 1 | - | 1 | 1 | - | 1 |

* Ig-λ datasets were used in the searches but in both cases the average and maximum scores were significantly lower compared to Ig-κ. Therefore Ig-λ sequences were not considered further.

**Table S3.** CryoEM data collection information

| | BG505 SOSIP + Rh.33104 mAb.1 + RM20A3 | BG505 SOSIP + Rh.33172 mAb.1 + RM19R |
|---|---|---|
| **Microscope** | Titan Krios | Titan Krios |
| **Voltage (kV)** | 300 | 300 |
| **Detector** | Gatan K2 Summit | Gatan K2 Summit |
| **Recording mode** | Counting | Counting |
| **Magnification** | 29,000 X | 29,000 X |
| **Movie micrograph pixel size** | 1.03 | 1.03 |
| **Dose rate ($e^{-}$/Å$^2$/s)** | 4.70 | 4.70 |
| **No. of frames per movie micrograph** | 38 | 38 |
| **Frame exposure time (ms)** | 250 | 250 |
| **Movie micrograph exposure time (s)** | 9.5 | 9.5 |
| **Total dose ($e^{-}$/Å$^2$)** | 44.7 | 44.7 |
| **Under focus range (µm)** | 0.8 – 1.6 | 0.7 – 1.6 |
| **Number of movie micrographs** | 1022 | 2050 |

**Table S4.** Model building and refinement information

| | BG505 SOSIP + Rh.4O9 pAbC-1 | BG505 SOSIP + Rh.33104 mAb.1 + RM20A3 | BG505 SOSIP + Rh.33172 mAb.1 + RM19R |
|---|---|---|---|
| **EMDB ID** | EMD-23779 | EMD-23780 | EMD-23801 |
| **Map Resolution (Å)** | 3.6 | 3.3 | 3.5 |
| **Map Symmetry** | C1 | C3 | C1 |
| **PDB ID** | 7MDT | 7MDU | 7MEP |
| **Residues** | 1970 | 1051 | 2686 |
| **Amino-acids** | 1892 | 1018 | 2596 |
| **Carbohydrates** | 78 | 33 | 90 |
| **RMSD Bonds** | 0.022 | 0.023 | 0.023 |
| **RMSD Angles** | 1.771 | 1.711 | 1.696 |
| **Ramachandran** | | | |
| **Outliers (%)** | 0.00 | 0.00 | 0.00 |
| **Allowed (%)** | 2.49 | 2.10 | 1.49 |
| **Favored (%)** | 97.51 | 97.90 | 98.51 |
| **Rotamer outliers** | 0.12 | 0.00 | 0.00 |
| **Clash score** | 0.63 | 1.04 | 0.89 |
| **Molprobity score** | 0.81 | 0.83 | 0.77 |
| **EMRinger score** | 3.38 | 4.32 | 3.66 |

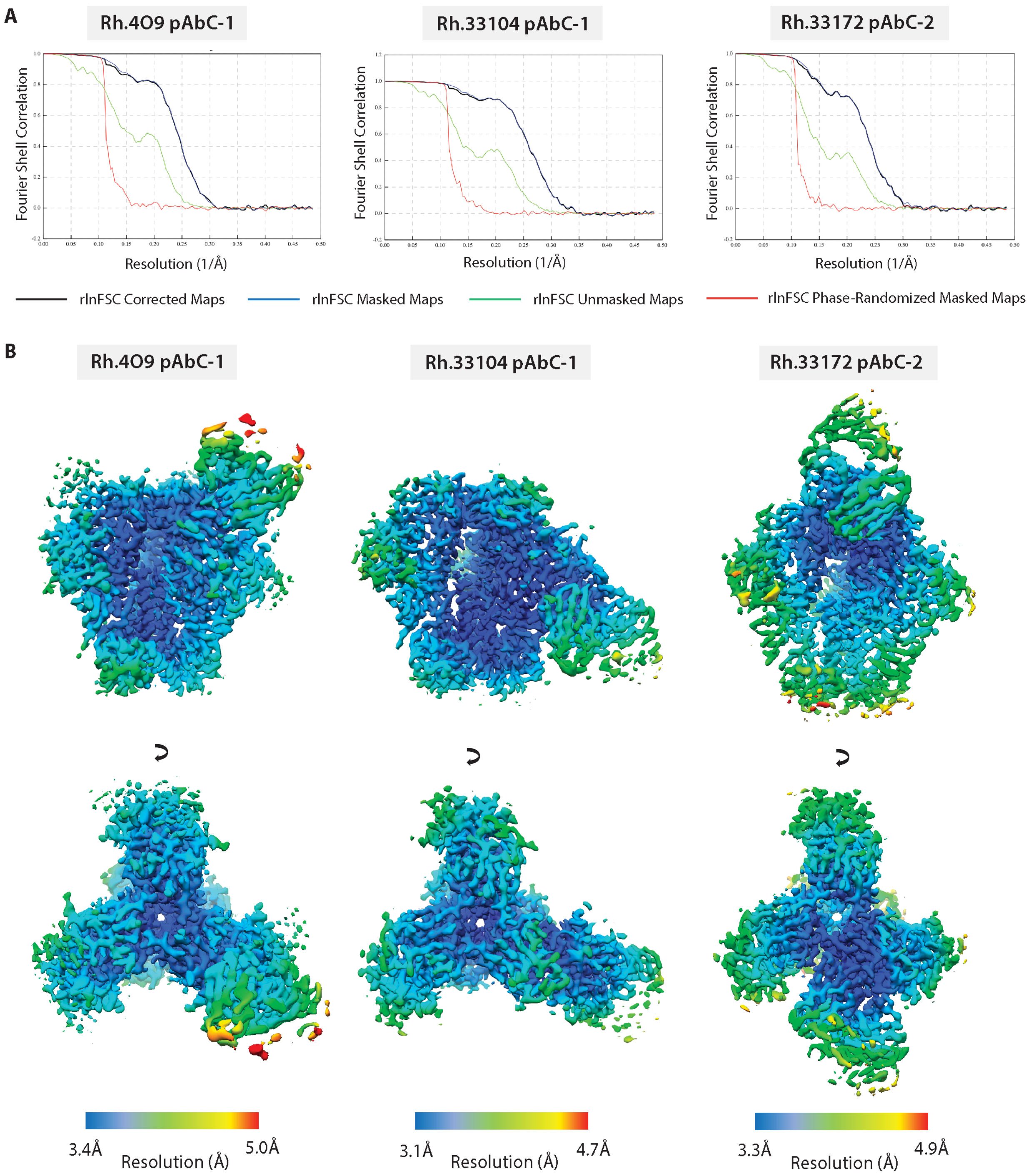


**Figure S1.** **[A]** Fourier shell correlation curves for trimer-pAbC complexes obtained using cryoEMPEM. **[B]** Local resolution plots for the trimer-pAbC complexes under investigation. The data for Rh.33172 pAbC-2 and Rh.33104 pAbC-1 were adapted from previously published work (*1*).

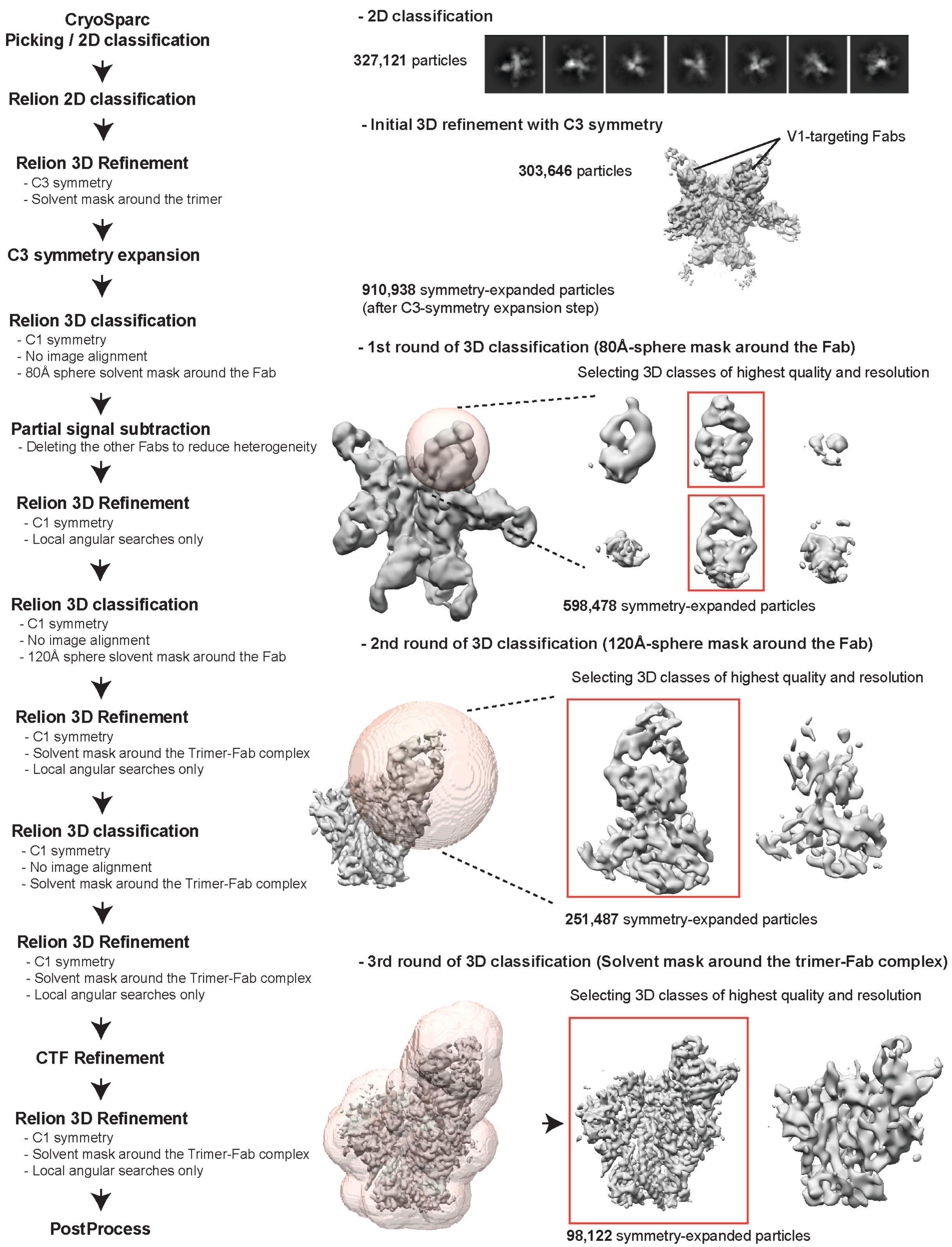


**Figure S2.** Schematic representation of the data processing workflow for Rh.4O9 cryoEMPEM data. Intermediate results and particle count data are shown on the right.

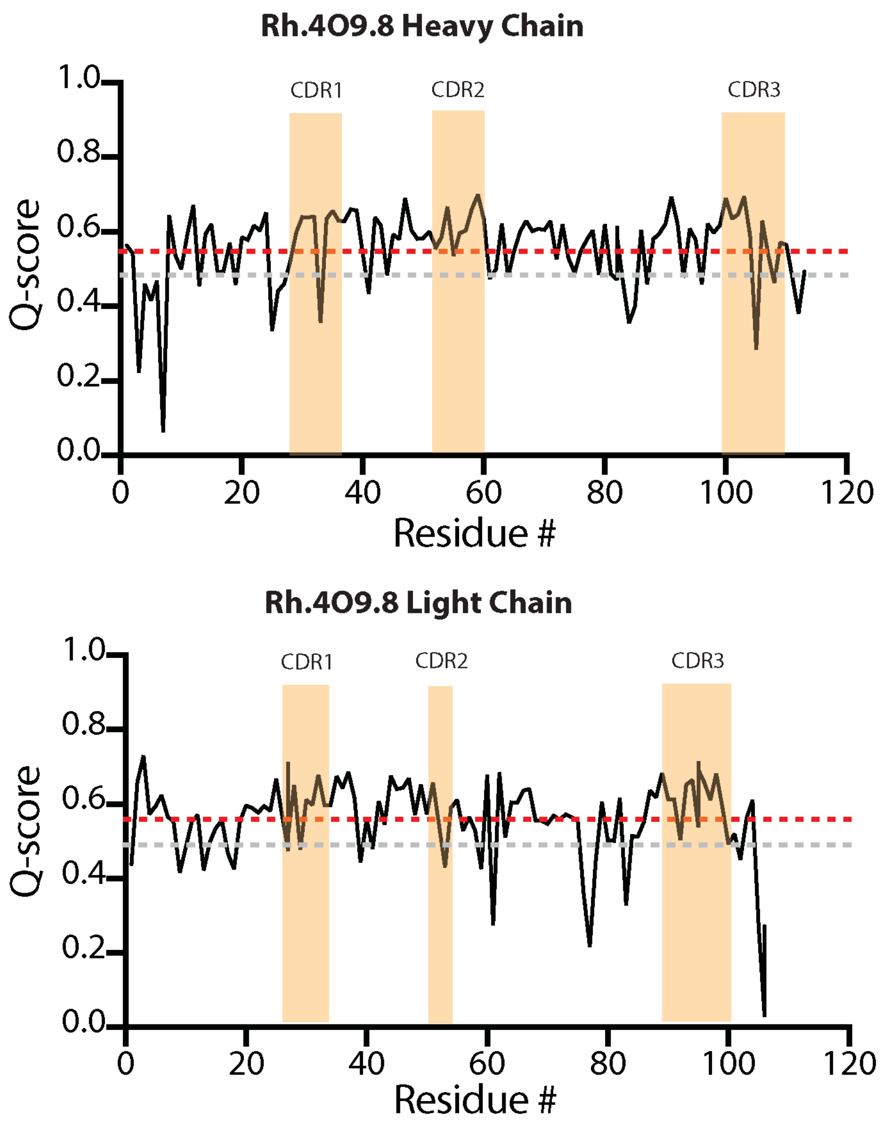


**Figure S3.** Analysis of the model-to-map fit for the Rh.4O9.8 antibody and the Rh.4O9 pAbC-1 cryoEMPEM map. Per-residue Q-score values for the heavy (top) and light (bottom) chains of the Rh.4O9. Average Q-score for each chain is displayed as dotted red line. The expected average Q-score for a map of 3.6 Å resolution is shown as a dotted gray line. CDRs are represented in orange.

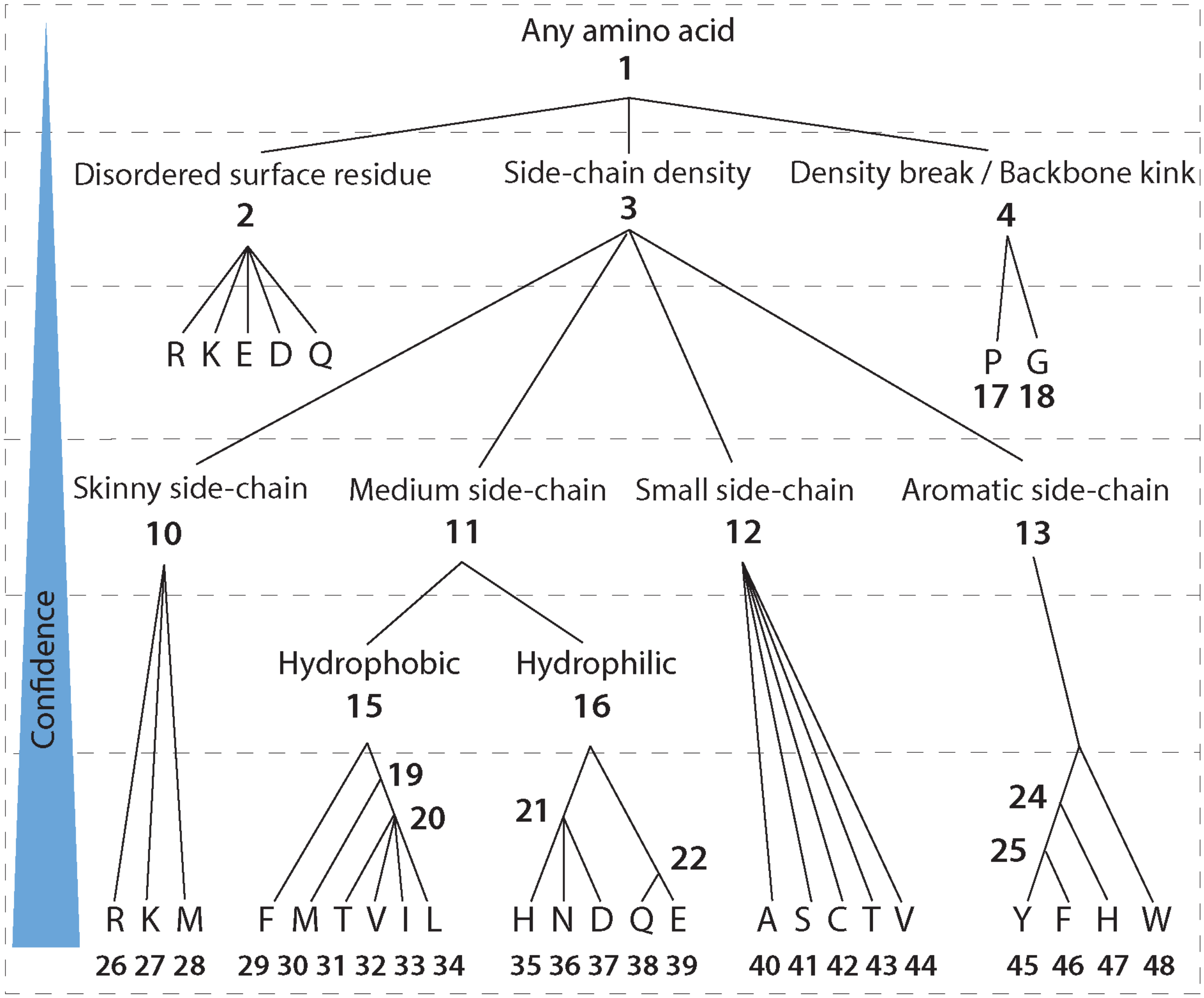


**Figure S4.** Amino acid assignment tree with numerical category identifiers. Amino acids that appear more than once under different codes (for example, Methionine can be represented as category 28 or 30) are treated as redundant entries in the search.

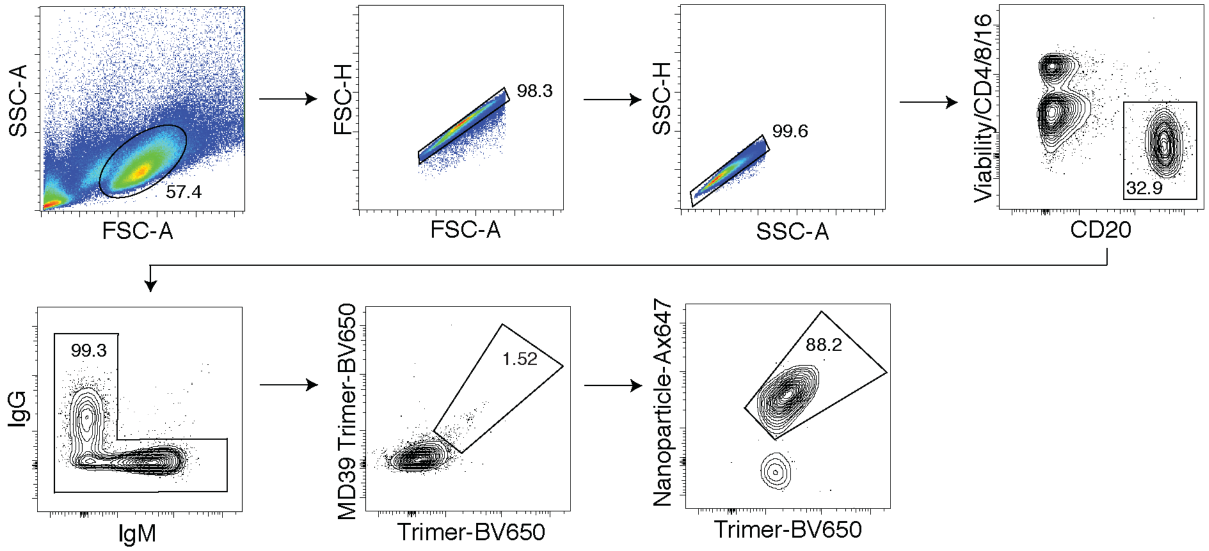


**Figure S5.** Overview of the gating strategy used for B-cell sorting. More detailed explanations are provided in the methods section.

**A**

**Rh.33104 pAbC-1 Light Chain**
**(residues 94-102)**

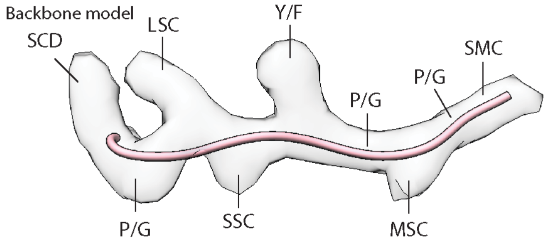


| Prediction | SCD | P/G | LSC | SSC | Y/F | P/G | MSC | P/G | SMC |
|---|---|---|---|---|---|---|---|---|---|
| Matching | + | + | + | + | + | + | + | + | + |
| Best Match | Y | P | R | T | F | G | Q | G | T |

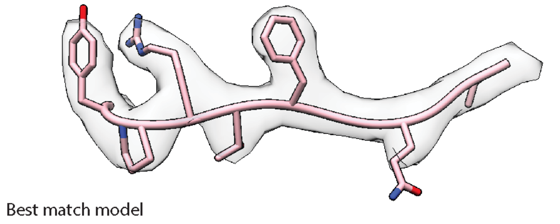


**Rh.33104 pAbC-1 Heavy Chain**
**(residues 102-110)**

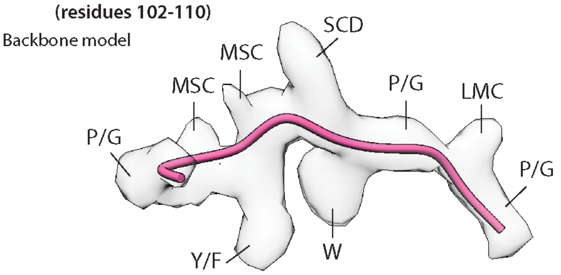


| Prediction | P/G | MSC | Y/F | MSC | SCD | W | P/G | LMC | P/G |
|---|---|---|---|---|---|---|---|---|---|
| Matching | - | + | + | + | + | + | + | + | + |
| Best Match | S | N | F | D | F | W | G | Q | G |

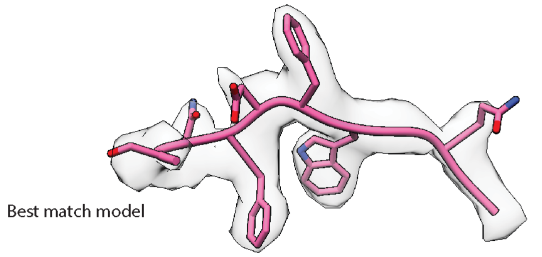


**B**

**Rh.33104 pAbC-1 Light Chain (kappa)**

| Segment | Sequence and matching | Matches | Score |
|---|---|---|---|
| FR1 | D I Q M T Q S P S S L S A S V G D T V T T T C R A S<br>+ + + + + + + + + + + + + + + + + + + + + + + + + + | 26/26 | 5.25 |
| CDR1 | Q D I S N D<br>+ + + + + + | 6/6 | 0.85 |
| FR2 | L A W Y Q Q K P G K A P K P L L Y<br>+ + + + + - + + + + + + + - + + + | 15/17 | 5.40 |
| CDR2 | Y A S<br>+ + + | 3/3 | 0.65 |
| FR3 | N L E S G V P S M F S G S G S G T D F T L T I S S L Q P E D F A S Y F C<br>+ + - + + + + + + + + + + + + + + + + + + + + + + + + + + + + + + + + + | 35/36 | 9.70 |
| CDR3 | Q Q Y N S Y P R T<br>+ + + + + + + + + | 9/9 | 1.89 |
| | Total | 94/97 | 23.74 |

**Rh.33104 pAbC-1 Heavy Chain**

| Segment | Sequence and matching | Matches | Score |
|---|---|---|---|
| FR1 | Q V Q L Q E S G P G L V K P S E T L S L T C A V S<br>- + - + - - - + + - - + + + + - + + + + + + + + - | 16/25 | 2.81 |
| CDR1 | G G S F S G Y S<br>+ + + + + + + - | 7/8 | 1.67 |
| FR2 | W G W I R Q P P G K G L E W I G S<br>+ + + + + + + + + + + + + + + + - | 16/17 | 7.32 |
| CDR2 | I I G R T G S T<br>+ + - + + + + + | 7/8 | 1.49 |
| FR3 | A Y N P S L T S R V T I S R D T S N N Q F S L K L T S L T A A D T A V Y Y C<br>+ + + + + + + + + + + + + + + - + + + + + + + + + + + + - + + - + + + + + + | 35/38 | 7.31 |
| CDR3 | A R Q Q S N F D F<br>+ + + + - + + + + | 8/9 | 1.42 |
| | Total | 89/105 | 22.02 |

**Figure S6. [A]** Example of the amino-acid assignment for Rh.33104 pAbC-1. For light chain (left), residues 94-102 and the corresponding area of the map are shown. For heavy chain (right), residues 102-110 and the corresponding area of the map are shown. Models are displayed in pink and maps are represented as transparent light-gray surface. **[B]** Best sequence matches for Rh.33104 pAbC-1 light chain (top) and heavy chain (bottom). Matching to assignments at each position is shown (+/-). Overall agreement to predictions and scoring data are shown on the right.

**A**

**Rh.33172 pAbC-2 Light Chain**
**(residues 32-40)**

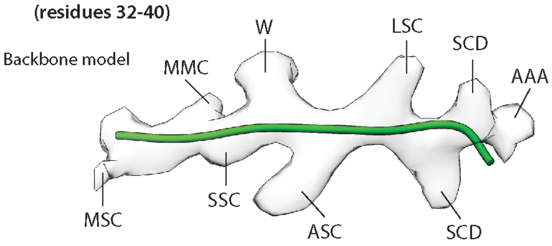


| Prediction | MSC | MMC | SSC | W | ASC | LSC | SCD | SCD | AAA |
|---|---|---|---|---|---|---|---|---|---|
| Matching | + | + | + | + | + | - | + | + | + |
| Best Match | Q | L | C | W | Y | Q | L | K | P |

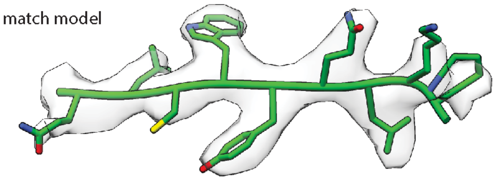


**Rh.33172 pAbC-2 Heavy Chain**
**(residues 107-115)**

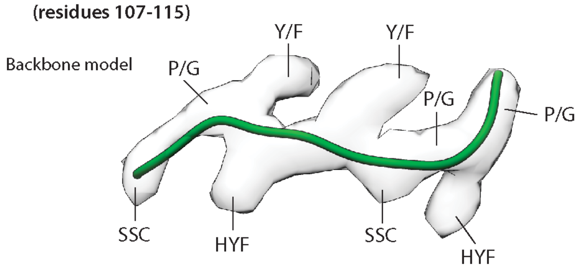


| Prediction | SSC | P/G | Y/F | HYF | Y/F | SSC | P/G | HYF | P/G |
|---|---|---|---|---|---|---|---|---|---|
| Matching | - | + | + | + | + | + | + | + | + |
| Best Match | D | G | Y | H | Y | T | G | Y | G |

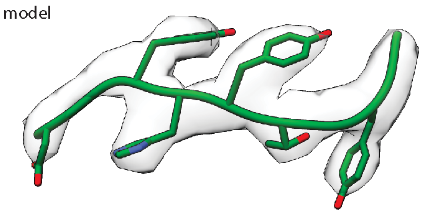


**B**

**Rh.33172 pAbC-2 Light Chain (kappa)**

| Segment | Sequence and matching | Matches | Score |
|---|---|---|---|
| FR1 | D I Q M T Q S P S S L S A S I G D R V T V T C R A S<br>+ - + - + - - + + + - + + - + + - + + + + - + + + + | 18/26 | 3.12 |
| CDR1 | Q G I N M Q<br>+ + + + + + | 6/6 | 1.03 |
| FR2 | L C W Y Q L K P G K A P T L L I Y<br>+ + + + - + + + + + + - + + + + + | 15/17 | 3.32 |
| CDR2 | G T S<br>+ + + | 3/3 | 0.90 |
| FR3 | G L Q T G V S S R F S G S G S G T N F T L T I S S L Q P E D V A T Y Y C<br>- + + + + + - + + + + + + - - + + - + + + + + + + + + + + + + - + + + + | 30/36 | 5.22 |
| CDR3 | Q Q D Y T T P F T<br>+ + + + + + - + + | 8/9 | 1.03 |
| | | **Total** | **80/97** | **14.62** |

**Rh.33172 pAbC-2 Heavy Chain**

| Segment | Sequence and matching | Matches | Score |
|---|---|---|---|
| FR1 | Q V Q L V Q S G A E V K M P G T S V K L S C K T S<br>+ + + + + - + + + + + - + + + + + + + + + + + + + | 23/25 | 3.94 |
| CDR1 | G Y T F T S Y N<br>+ + + + + + + - | 7/8 | 1.62 |
| FR2 | I N W V R Q A P G Q A L E W M G W<br>+ - + + + + + - + + + + + + + + + | 15/17 | 4.88 |
| CDR2 | I N P N N G T T<br>+ + + + + + + + | 8/8 | 1.16 |
| FR3 | D Y A Q K F Q G R V T M T R D T S T T T A Y M Q L N S L R S E D T A V Y Y C<br>+ + + - + + + + + + + + + - - + - + + + + + + + + - + + + + + + + + + + + + | 33/38 | 7.18 |
| CDR3 | A R A R G G Y E D D D G Y H Y T G Y G L D<br>+ + + - - + + + + - - + + + + + + + + + + | 17/21 | 5.37 |
| | **Total** | **103/117** | **24.15** |

**Figure S7. [A]** Example of the amino-acid assignment for Rh.33172 pAbC-2. For light chain (left), residues 32-40 and the corresponding area of the map are shown. For heavy chain (right), residues 107-115 and the corresponding area of the map are shown. Models are displayed in green and maps are represented as transparent light-gray surface. **[B]** Best sequence matches for Rh.33172 pAbC-2 light chain (top) and heavy chain (bottom). Matching to assignments at each position is shown (+/-). Overall agreement to predictions and scoring data are shown on the right.

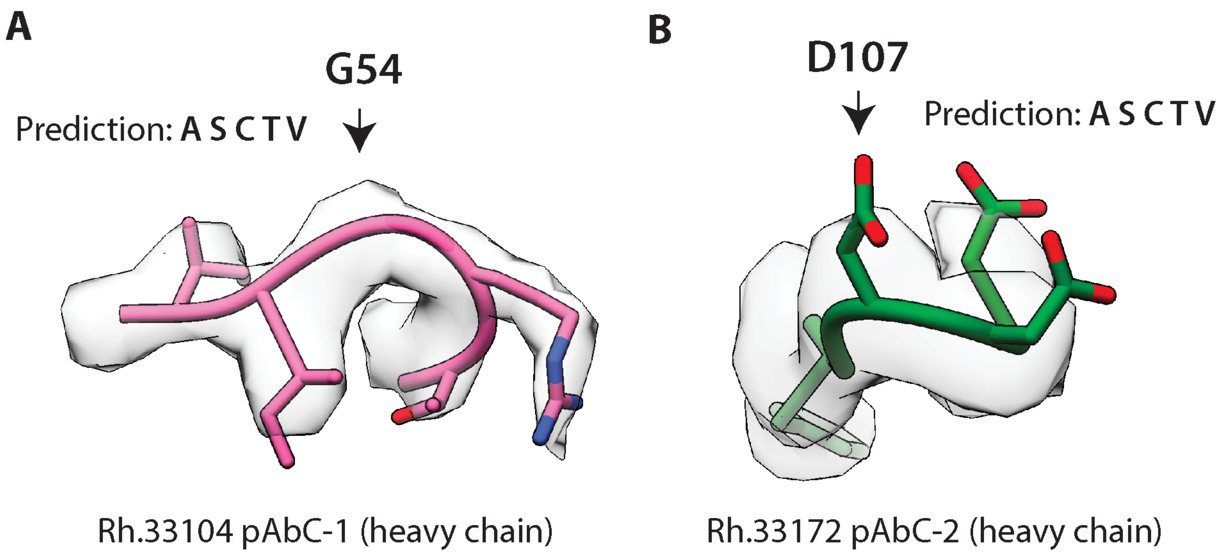


**Figure S8.** Examples of the most common mismatches observed in the Rh.33104 (pink) and Rh.33172 searches (green). **[A]** Small side-chain category (amino acids: A, S, C, T, V) was predicted based on structural data but P/G category (amino acids: P and G) was in the sequence. **[B]** Small side-chain category (amino acids: A, S, C, T, V) was predicted based on structural data but medium side-chain LMC category (amino acids: H, N, D, Q, E) was in the sequence. Map segments are represented as transparent light-gray surface.

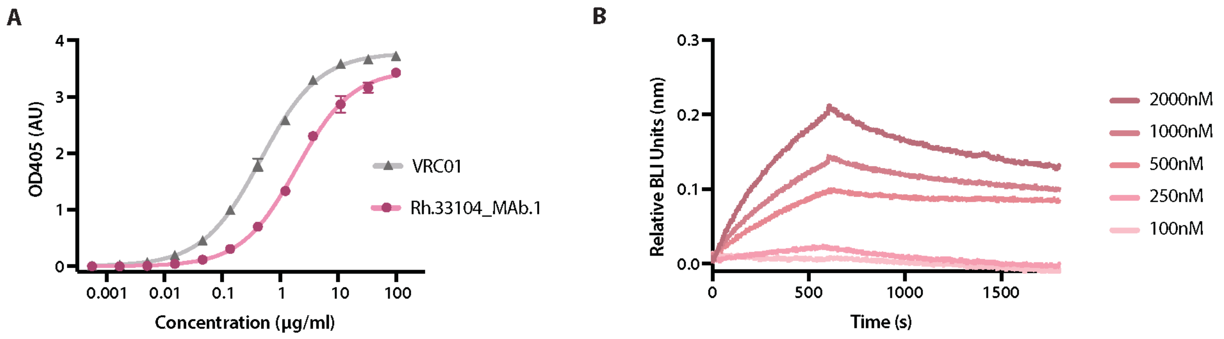


| | Rh.33104 mAb.1 | VRC01 |
|---|---|---|
| EC50 (µg/ml) | 1.93 | 0.46 |

| | kon (1/M*s) | koff (1/s) | Kd (M) |
|---|---|---|---|
| Rh.33104 mAb.1 | $5.29 * 10^2$ | $4.76 * 10^{-4}$ | $(8.9 \pm 0.6) * 10^{-7}$ |

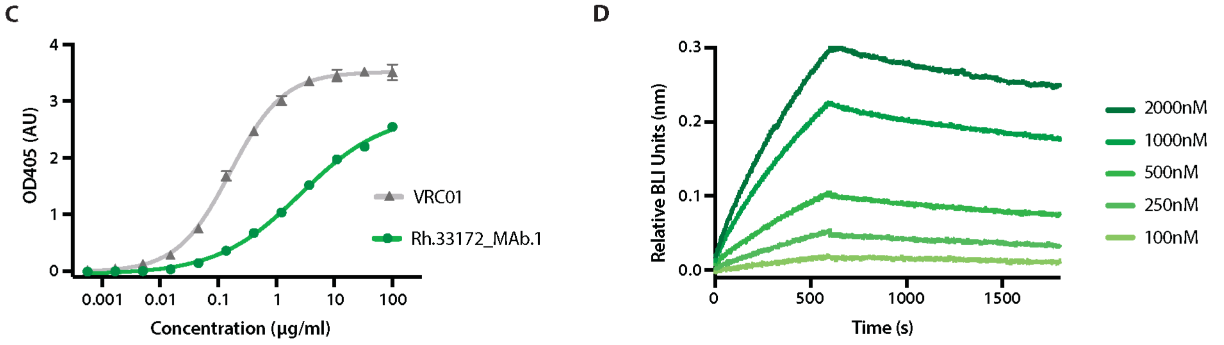


| | Rh.33172 mAb.1 | VRC01 |
|---|---|---|
| EC50 (µg/ml) | 2.64 | 0.16 |

| | kon (1/M*s) | koff (1/s) | Kd (M) |
|---|---|---|---|
| Rh.33172 mAb.1 | $5.03 * 10^2$ | $9.08 * 10^{-5}$ | $(1.80 \pm 0.03) * 10^{-7}$ |

**Figure S9.** Binding data for Rh.33104 mAb.1 and Rh.33172 mAb.1. Sandwich ELISA was used to quantify the interaction of BG505 SOSIP to the IgG versions of Rh.33104 mAb.1 **[A]** and Rh.33172 mAb.1 **[C]**. VRC01 IgG was used as a positive control and reference. $EC_{50}$ values are in the tables below the corresponding graphs. BLI was used to determine the kinetic binding parameters of the interaction between BG505 SOSIP and the Fab versions of Rh.33104 mAb.1 **[B]** and Rh.33172 mAb.1 **[D]**. Antigen concentrations used to generate each binding curve are illustrated on the right in panels **[B]** and **[D]**, while the corresponding kinetic parameters are presented in the table below each graph.

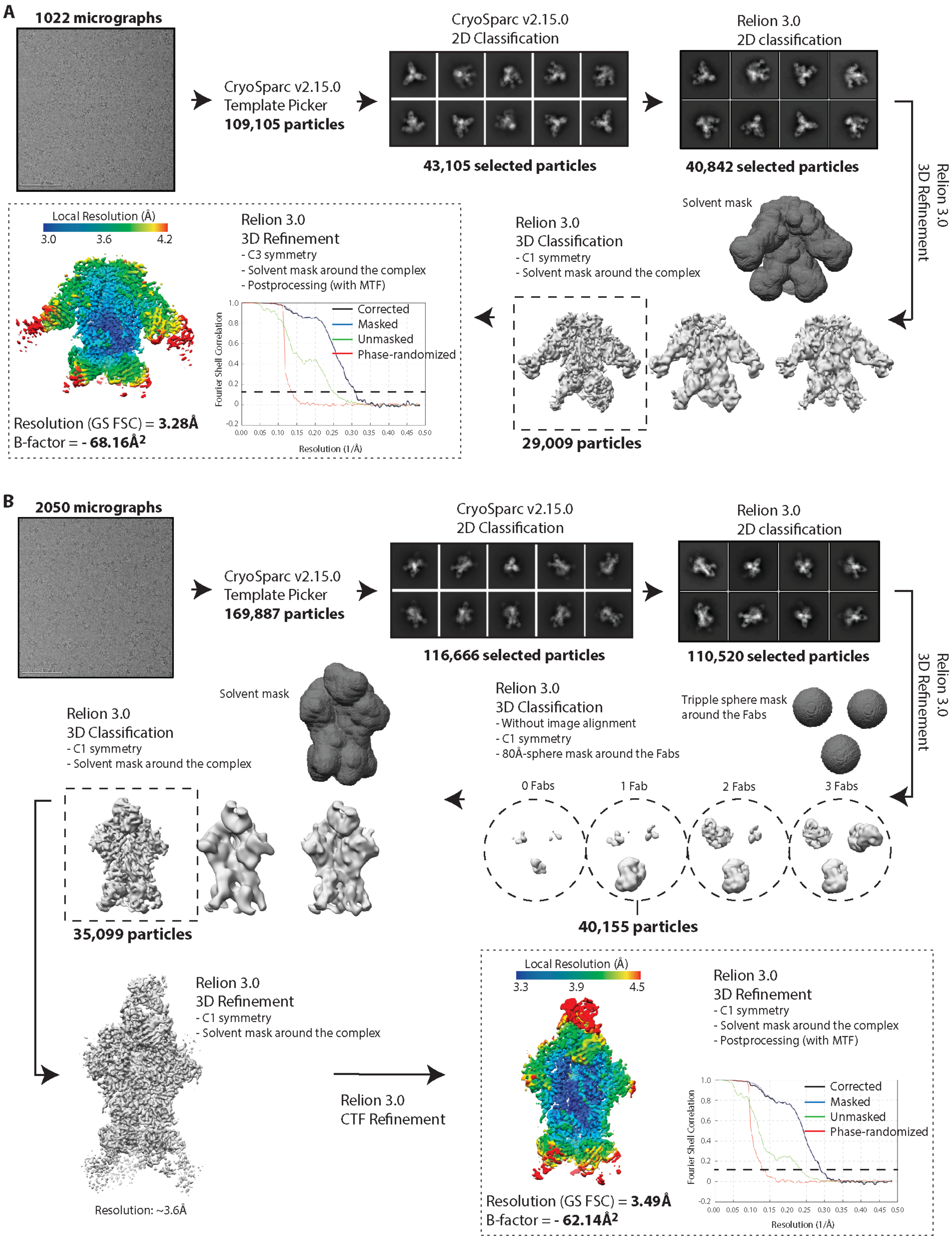


**Figure S10.** Schematic representation of the data processing workflow for cryoEM data with relevant statistics. The samples were **[A]** BG505 SOSIP complexed with Rh.33104 mAb.1 and RM20A3 and **[B]** BG505 SOSIP complexed with Rh.33172 mAb.1 and RM19R

#### Supplementary Material References
